# Establishing the Median Infectious Dose (ID_50_) and Characterizing the Clinical Manifestations of Mouse, Rat, Cow, and Human *Corynebacterium bovis* Isolates in Select Immunocompromised Mouse Strains

**DOI:** 10.1101/2022.11.10.515978

**Authors:** Gerardo Mendoza, Christopher Cheleuitte-Nieves, Kvin Lertpiriyapong, Juliette R.K. Wipf, Rodolfo J. Ricart Arbona, Ileana C. Miranda, Neil S. Lipman

## Abstract

*Corynebacterium bovis* (Cb), the etiology of hyperkeratotic dermatitis in various immunocompromised mouse strains, significantly impacts research in which infected mice are used. Although Cb has been isolated from a variety of species, including mice, rats, cows, and humans, little is known about the differences in the infectivity and clinical disease in mice associated with unique isolates. The infectious dose yielding colonization of 50% of the exposed population (ID_50_) and any associated clinical disease was determined for mouse (n=3), rat (n=1), cow (n=1), and human (n=2) Cb isolates in athymic nude mice (Hsd:Athymic Nude-Foxn1^nu^). The same investigations were undertaken comparing 2 of these murine isolates in 2 furred immunocompromised mouse strains (NSG [NOD.Cg-Prkdc^scid^Il2rg^tm1Wjl^/Sz] and NSG-S [NOD.Cg-Prkdc^scid^Il2rgt^m1Wjl^Tg(CMV-IL3,CSF2,KITLG)1Eav/MloySzJ]). To determine the ID_50_, mice (n=6/dose; 3 of each sex) were inoculated topically with 1 to 10^8^ bacteria (10-fold increments) of each Cb isolate. Mice were scored (0 to 5) daily based on the severity of clinical signs for 14 days. On day 7 and 14 post-inoculation (PI), buccal and dorsal skin swabs were evaluated by aerobic culture to determine infection status. The mouse isolates yielded a lower ID_50_ (58 to 1,000 bacteria) as compared to the bovine (6,460 to 7,498 bacteria) and rat (10,000 bacteria) Cb isolates. Mice were not colonized and disease did not result when inoculated with human isolates. Mouse isolates produced varying clinical disease severity in nude mice (max score/isolate: 0 to 5). Despite significant immunodeficiency, furred NSG and NSG-S mice required a considerably higher (1,000- to 3,000-fold) inoculum to become colonized as compared to athymic nude mice. Once colonized, clinically detectable hyperkeratosis did not develop in these strains until 18 to 22 days PI. In contrast, in athymic nude mice that developed clinically detectable disease, hyperkeratosis was observed 6 to 14 days PI. In conclusion, there are significant differences in Cb’s ID_50_, disease course, and severity between Cb isolates and among immunodeficient mouse strains.

## INTRODUCTION

*Corynebacterium*-associated hyperkeratosis (CAH) has been reported anecdotally as early as 1976 in athymic nude mice with subsequent global outbreaks described in the 1980’s and 1990’s.^11,30^ In 1998, *Corynebacterium bovis* (Cb), the causative agent of CAH, also known as scaly skin disease, was speciated using 16S rRNA sequence analysis.^14^ Cb, a Gram-positive, rod-shaped bacterium, causes opportunistic infections in immunocompromised mice including, but not limited to, the *Foxn1^nu−/−^* athymic nude (NU), NOD.Cg-Prkdc^scid^Il2rg^tm1Wjl^ (NSG), and Pkrdc^scid^ (SCID) strains.^11,31^ Additionally, the bacterium causes similar lesions in immunocompetent epidermal mutant *dep/dep* and hairless SKH1-Hr^hr^ mice, presumably due to sebaceous gland hyperplasia as well as altered follicle and hair shaft development.^11,26^ In nude and furred immunodeficient strains, clinical signs include diffuse hyperkeratotic dermatitis and alopecia with adherent white keratin flakes on the skin, respectively.^5,39^ The clinical syndrome in nude mice typically develops within 7-10 days post exposure and may include weight loss, dehydration and pruritus.^11,13,30^ In contrast, there are limited studies assessing Cb infection in furred immunocompromised mice and there are no published studies that describe the disease course in NSG mice, which are commonly used and frequently infected with Cb.^31^

In mice with CAH, the epidermis is hyperplastic, displaying acanthosis and orthokeratotic hyperkeratosis, which may be accompanied by mild dermal mononuclear cell infiltrates.^35^ Lesion severity is thought to be multifactorial and may be exacerbated by experimental manipulation including administration of chemotherapeutics, surgery, and irradiation.^5^

Cb is spread by direct contact with contaminated keratin flakes, fecal-oral, and airborne transmission, on fomites (cages, gloves, bedding, work surface, and instruments), or through contaminated biologics including cryopreserved tumors.^5,24,39^ Additionally, Cb has been isolated from biosafety cabinets and the respiratory tract of animal care staff.^6^ Introduction of Cb into a vivarium generally leads to widespread colonization of susceptible stocks and strains, with high prevalence of clinical disease in immunocompromised mice.^30^ Once infected, NU mice remain colonized despite antibiotic treatment and clinical resolution.^5,25^ Furthermore, the use of antibiotics in tumor models may confound research as their use can alter the tumor microenvironment and make them more resistant to future treatments including chemotherapeutics.^22^ Cb has been shown to affect engraftment of chronic myelomonocytic leukemia in NSG-S mice and is thought to alter the immune profile of infected animals.^38^

Once CAH is suspected, diagnosis is confirmed by bacterial isolation or PCR.^5,23^ The bacterium can be reliably isolated from a skin or buccal swab with high sensitivity reaching 100% and 94% in NU mice, respectively.^5^ A thorough understanding of Cb’s infectivity, pathogenicity, and epidemiology is critical to developing an effective strategy to prevent its introduction into susceptible colonies. Cb has been isolated from humans and various animal species and genome-wide analyses have shed light on the differences in the genetic composition of these isolates and uncovered putative virulence factors and pathogenicity islands.^9^ Based on their genetic sequence, the isolates clustered into 2 distinct groups: 1) rodent isolates; and 2) bovine and human isolates.^9^ However, relatively little is known as to whether these genetic differences influence Cb’s infectivity, pathogenicity, infectivity, or its inter-species transmission potential.

An important consideration in understanding the infectivity of a microorganism is to determine the median number of microbial units necessary to effectively colonize 50% of the exposed population, referred to as the median infectious dose (ID_50_).^2,7,28^ Characterizing infectivity can help determine if certain populations within the vivarium are more susceptible and biosecurity protocols can be developed based on these findings. Historically, 3 methods have been used to estimate the ID_50_ from dose response data, the Reed–Muench, Dragstedt–Behrens, and Spearman–Karber methods.^19,28,29,37^ More recently, probit, logistic regression, exponential, and approximate Beta Poisson models have been used for quantitative microbial risk assessment allowing the probability of infection to be estimated based on any dose.^19^

This study was undertaken to determine the ID_50_ and dose response curves, as well as to characterize the clinical disease associated with Cb isolates cultured from mice, rat, bovine, and humans. Additionally, the ID_50_ and associated clinical disease of 2 Cb mouse isolates were determined and compared in NU, NSG and NSG-S mice.

## MATERIALS AND METHODS

### Study Design

The ID_50_ of 9 Cb isolates from various species and institutions (Table 1) were determined in NU mice. The ID_50_ was also determined in the NSG strain for isolates 7984 and 13-1426, and the NSG-S strain for isolate 7894. To establish the ID_50_ in NU mice, up to 8 doses (1 to 10^8^ in 10-fold increments) of each isolate in mid-log growth phase were administered topically to 6 mice (3 of each sex). For each isolate, 2 mice (1 of each sex) served as uninfected controls. Following infection, each mouse was singly housed. Post-inoculation (PI), mice were monitored daily for 14 days and scored based on clinical signs. On day 7 and 14 PI, mice were weighed, and buccal and dorsal skin swabs were collected to assess Cb colonization by aerobic culture. Mice were euthanized by CO_2_ overdose on day 14 PI, the body and splenic weights were determined, a postmortem examination was performed, and macroscopic changes documented, and full thickness, 6 mm diameter, skin punch biopsy samples were collected from the nuchal crest, and the dorsal lumbar region for histopathology. Additional biopsy samples were taken from lesions, if present. To determine the clinical presentation of Cb in NSG mice, 4 female NSG mice were each inoculated with Cb isolate 7984 or 13-1426 (10^8^ bacteria/mouse). Following inoculation, 2 mice receiving the same inoculum were cohoused with 2 naïve NSG mice and monitored daily for signs of disease for 60 days. The animals were weighed, and dorsal skin swabs were collected from each mouse every 14 days PI to determine the presence and estimated quantity of Cb present. Mice were euthanized on day 60 PI.

**Table 1.** Characteristics of the *Corynebacterium bovis* isolates utilized including the species from which the isolate was cultured, isolation site, institutional source, and the number of putative virulence factors reported.^9^

| Isolate ID | Host | Isolation Site | Institution | Putative virulence factors [#] |
| --- | --- | --- | --- | --- |
| 7894 | Athymic nude Mouse | Skin disease (hyperkeratosis and acanthosis) | MSK | 29 |
| CUAMC1 | Athymic nude Mouse | Skin disease (hyperkeratosis and acanthosis) | UC - Anschutz Medical Campus | 44 |
| 13-1426 | NSG mouse | Skin disease (hyperkeratosis and acanthosis) | Biopharma | 79 |
| 17-0240 | Athymic nude Mouse | Skin disease (hyperkeratosis and acanthosis) | MSK | 28 |
| Iso 24956 | Athymic nude mouse | Skin (no clinical disease) | JAX | Unknown |
| 16-2004 | Athymic nude Rat | Skin disease (dermatitis and ulcerative lesions) | MSK | 75 |
| 4826 | Bovine | Mastitis/milk | CDC SBRL | 33 |
| F6900 | Human, female | Wound | CDC SBRL | 19 |
| WCM 35 | Human, female | Eye | NYPH-WCM | Unknown |

Abbreviations: MSK, Memorial Sloan Kettering Cancer Center; UC, University of Colorado; JAX, The Jackson Laboratory; CDC SBRL, Centers for Disease Control and Prevention, Special Bacteriology Reference Laboratory; NYPH-WCM, New York Presbyterian Hospital - Weill Cornell Medicine.

### Animals

NU (Hsd:Athymic Nude-Foxn1^nu^; Envigo, Indianapolis, IN; *n*=414), NSG (NOD.Cg-Prkdc^scid^Il2rg^tm1Wjl^/Sz; The Jackson Laboratory [Jax], Bar Harbor, ME; *n*=100), and NSG-S (NOD.Cg-Prkdcscid Il2rgtm1Wjl Tg (CMV-IL3, CSF2, KITLG)1Eav/MloySzJ; Jax; *n*=50) mice were F1 generations bred in-house. Mice were free of the following agents: mouse hepatitis virus, Sendai virus, mouse parvovirus, minute virus of mice, murine norovirus, murine astrovirus- 2, pneumonia virus of mice, Theiler meningoencephalitis virus, epizootic diarrhea of infant mice (mouse rotavirus), ectromelia virus, reovirus type 3, lymphocytic choriomeningitis virus, K virus, mouse adenovirus 1 and 2, polyoma virus, murine cytomegalovirus, mouse thymic virus, Hantaan virus, murine astrovirus-2, *Mycoplasma pulmonis, Citrobacter rodentium, Salmonella spp., Filobacterium rodentium, Clostridium piliforme, Chlamydia muridarum, Corynebacterium bovis*, fur mites (*Myobia musculi, Myocoptes musculinus, and Radfordia affinis), pinworms (Syphacia spp. and Aspicularis tetraptera*), and *Encephalitozoon cuniculi*. A day prior to Cb inoculation, the dorsal thoracolumbar skin of the mice was swabbed and confirmed negative for Cb by culture.

### Husbandry and Housing

Pre-inoculation, mice were maintained in autoclaved, polysulfone individually ventilated cages (IVCs) with stainless-steel wire-bar lids and filter tops (no. 19, Thoren Caging Systems, Inc., Hazelton, PA) on autoclaved aspen chip bedding (PWI Industries, Quebec, Canada) at a density of no greater than 5 mice per cage. Post-inoculation, mice were singly housed in sterile, solid-bottom and top, gasketed and sealed, polysulfone, positive pressure, IVCs (Isocage, Allentown Caging Equipment Company, Inc., Allentown, NJ) on autoclaved aspen-chip bedding (PWI Industries Canada, Quebec, Canada). Pre- and post-inoculated animals were provided autoclaved feed (5KA1, PMI, St Louis, MO) and autoclaved acidified (HCl) reverse osmosis water (pH 2.5 to 2.8) *ad libitum*. Each cage was provided with a sterile bag constructed of Glatfelter paper containing 6 g of crinkled paper strips (EnviroPak, WF Fisher and Son, Branchburg, NJ) for enrichment. Each cage was ventilated with approximately 30 air changes per hour of HEPA-filtered air (filtration at cage level) and the rack effluent was exhausted directly into the building’s exhaust system. Cages containing mice used for ID_50_ determination were not changed following inoculation until the mouse was euthanized at 14 days PI to reduce the likelihood of cross contamination. Cages containing NSG mice used to determine the clinical presentation of Cb and all post-inoculation cages were changed every 2 weeks. Cages were changed and all animal manipulations were performed within a class II, type A2 biological safety cabinet (LabGard S602-500, Nuaire, Plymouth, MN). The animal holding room was ventilated with at least 10 air changes of 100% fresh air hourly and maintained at 72 ± 2 °F (21.5 ± 1 °C), relative humidity between 30% and 70%, and a 12:12 hour light:dark photoperiod. All animal use described in this investigation was approved by MSK’s IACUC and conducted in agreement with AALAS’ position statements on the Humane Care and Use of Laboratory Animals and Alleviating Pain and Distress in Laboratory Animals. The animal care and use program is AAALAC International-accredited and operates in accordance with the recommendations provided in the Guide for the Care and Use of Laboratory Animals (8^th^ edition).^20^

### Bacterial culture methods

Cb isolates from skin and buccal swabs or frozen stocks were grown on trypticase soy agar supplemented with 5% sheep blood (BBL™ TSA II 5% SB, Becton, Dickinson and Company, Sparks, MD, USA) at 37°C with 5% CO_2_ for 48 hours as previously described.^5,10^ To determine the growth kinetics of each Cb isolate, 3 growth curves were established for each isolate, except isolate 7894 because its’ growth curve was previously established.^10^ Frozen stocks (Microbank TM, Pro-lab Diagnostics, Rounds Rock, TX) of each Cb isolate were grown as described above. Three colonies of each isolate were transferred to a 15-mL conical tube (Corning™ Falcon™, Fischer Scientific, Hampton, NH) containing 1.5 ml of brain–heart infusion broth ([BHI]; Becton Dickinson, Franklin Lakes, NJ) supplemented with 0.1% Tween 80 (VWR Chemicals, Solon, OH) and were incubated at 37°C with 5% CO_2,_ lying flat on a titer plate shaker (4625 Titer Shaker, Thermo Scientific, Waltham, MA) at 200 RPM. After 12h, an aliquot was collected and diluted 1:100 with fresh broth and incubated as described above. At least 5 aliquots were removed from the broth culture at intervals of 6 or 12 h for up to 96 h. Each aliquot was diluted 1:10 in sterile saline (5mL Saline Normal; Becton, Dickinson and Company, Sparks, MD, USA). The McFarland density was determined for each diluted aliquot using a densitometer (Den-1, Grantin Instruments, Shepreth, Cambridgeshire, UK) which measured the absorbance of light at a wavelength of 565±15 nanometers and provided a numeric value corresponding to the McFarland interpretation. If required, an additional dilution was made to obtain a reading within the densitometer’s dynamic range. A Gram stain was performed on dilutions to confirm the culture remained contaminate free. McFarland values were plotted, and the mid-log was determined on GraphPad Prism using a nonlinear sigmoidal 4PL best fit analysis based on the 50% end point. To determine the approximate cfu/ml of Cb cultures in exponential phase of growth, Cb isolates were grown in triplicate as described above. Aliquots were taken at time points corresponding to mid-exponential growth and enumeration of bacteria was performed by 10-fold serial dilution and plating.

### Animal inoculation

The bacterial inoculum collected from the culture during the mid-log of the exponential growth phase was titrated and serially diluted in BHI broth to obtain 1 to 10^8^ +/− 15% bacteria/50 μl in 10-fold increments for all isolates except 7894, CUAMC1, and F6900, for which a maximum inoculum of 10^4^, 10^6^, and 10^6^ bacteria, respectively, was administered. These isolates were the initial 3 isolates evaluated and the dose range used for the earliest ID_50_ determinations was based on a pilot study conducted with isolate 7894 in which all inoculated mice were colonized with inocula below 10^3^ bacteria. The dose range was expanded for CUAMC1 and F6900, the next isolates evaluated, to 10^6^ bacteria and expanded further to 10^8^ bacteria for all subsequent isolates evaluated when a 10^6^ inoculum was determined to be insufficient necessitating the administration of the maximum inocula used of 10^8^ bacteria/mouse. Cb inocula were placed in sterile 15 ml polypropylene centrifuge tubes (Fisher Scientific, Waltman, MA), transferred to the vivarium on ice and animal inoculation was performed within an hour of preparation. After inoculation, the concentration of each inoculum was confirmed and enumerated by plate count.

Animal procedures, including inoculation, culture, and euthanasia, and cage change were performed in a class II, type A2 biologic safety cabinet ([BSC]; LabGard S602-500, Nuaire, Plymouth, MN) using sterile materials and aseptic methods by personnel donning chlorine dioxide disinfectant-wetted (Clidox [1:4:1], Pharmacal Laboratories, Naugatuck, CT) plastic sleeves and disposable gloves. Inoculation and handling of mice post-inoculation were performed in order of increasing inoculum dose. Each cage containing an experimentally naïve animal was removed from the rack and all sides of the cage were liberally sprayed with disinfectant and placed in the BSC. Each cage was opened for less than 2 minutes. Bacterial inoculation was performed with the mouse restrained with the handler’s non-dominant hand by gently grasping the tail base and the animal’s hindquarters were elevated slightly while the animal grasped the wire bar lid with its forelimbs. The lid was positioned to allow the restrained animal to remain in the horizontal plane. Using the dominant hand, the bacterial inoculum was applied directly to the skin using a sterile filter micropipette (P200N, Marshal Scientific, Hampton, NH) along the dorsal midline starting at the nuchal crest proceeding to the tail, a distance of ~ 2 cm. In furred mice, the fur was parted along the midline before applying the inoculum. A 50 μl bacterial suspension was evenly distributed along the dorsal midline at 4-5 sites dispensing 10-15 μL per site using a sterile filter 200 uL pipette tip (Universal Pipette Tips, Avantor, Bridgeport, NJ). This method ensured that the bacterial suspension did not flow off the animal. The cage was closed and sprayed thoroughly with disinfectant prior to returning it to the rack. All interior surfaces of the BSC were sprayed with disinfectant and a new pair of disinfectant wetted gloves and sleeves were donned between each cage. A new cage was placed in the BSC as previously described and the previous steps repeated until all inoculations (0 to 10^8^ bacteria/mouse) were completed. Animals in the control group were inoculated with an equivalent amount of bacterial-free BHI broth.

### Clinical scoring system

NU, NSG, and NSG-S mice used for ID_50_ determination and NSG mice used to assess clinical presentation were monitored daily for 14 or 60 days, respectively. Animals were evaluated cage side and scored using the system provided in Table 2 & 3.

**Table 2.** Clinical scoring system for nude mice. Animals were graded 0 to 5 based on clinical signs representing mild, moderate, and severe disease.

| Grade | Clinical Signs |
| --- | --- |
| 0 | No clinical signs observed. |
| 1 | Mild amount of flakes/scales adhered to the skin and/or mild hunched posture with no erythema. |
| 2 | Mild amount of flakes/scales adhered to skin and/or mild hunched posture and mild erythema associated with flaking/scaly lesions. |
| 3 | Moderate erythema and/or moderate amount of white flakes adherent to the skin, or minimal white flakes on 3 or more additional locations affected including the ventrum, muzzle, cheeks, crown, and limbs. |
| 4 | Moderate erythema and/or moderate amount of white flakes adherent to the skin and moderate hunched posture. |
| 5 | Severe amount of flakes/scales adhered to skin (thick layer), and/or severe erythema, and/or severe hunched posture. |

**Table 3.** Clinical scoring system for furred mice, e.g., NSG and NSG-S. Animals were graded 0 to 5 based on clinical signs representing mild, moderate, and severe disease.

| Grade | Clinical Signs |
| --- | --- |
| 0 | No clinical signs observed. |
| 1 | Ear base vessel dilation OR periorbital swelling and/or rough or unkempt hair coat. |
| 2 | Auricular hyperemia and periorbital swelling and/or rough or unkempt hair coat. |
| 3 | Periorbital swelling and/or auricular hyperemia and/or facial alopecia. |
| 4 | Severe periocular erythema/alopecia and/or a severe amount of auricular white flaking and truncal alopecia. |
| 5 | Severe periocular erythema/alopecia and/or a severe amount of auricular white flaking with thickened skin. |

### Post-inoculation bacterial sampling

Infection was assessed by aerobic culture of the skin and buccal region weekly for 2 weeks PI and the skin only once every 14 days PI in the 60-day NSG clinical presentation study. These sites were selected based on prior studies.^5^ All mice were sampled in order from lowest to highest inoculum. To prevent cross contamination within each dose group, animals with lower clinical scores were swabbed first. If animals had a similar score, cages were selected at random. All cages were handled aseptically as described in the animal inoculation section. Each cage was opened for no longer than 5 minutes. The animal was gently grasped by the base of the tail allowing all four limbs to grasp the wire bar lid. A sterile culture swab (Remel Bactiswab, Fisher Scientific, Waltman, MA), held perpendicular to the animal, was swabbed along the dorsal midline from the base of the tail cranially to the nuchal crest and caudally back to the tail base rotating the swab as it was advanced. For the caudal buccal swab, a sterile cotton tipped applicator (McKesson #24-106-2S, Irving, TX) was used as previously described.^5^ Briefly, the animal was restrained with the non-dominant hand grasping the animal using a three-finger technique. The animal was then weighed and returned to its cage. The scale used to ascertain body weight was wiped down with chlorine dioxide disinfectant and allowed to dry for 5 minutes prior to use. The cage was closed, sprayed, and wiped down with disinfectant and returned to the rack. The BSC was cleaned as previously described, a new pair of disinfected gloves was donned, and the next cage was processed after waiting 5 minutes.

### Confirmation of infection

The skin (PI days −1, 7, 14, and every 14 days for the NSG disease presentation study) and buccal swabs (PI days 7 and 14) were each streaked using a 4-quadrant pattern onto TSA II 5% SB agar plates (Becton Dickinson).^3^ The plates were incubated for 72 h at 37°C with 5% CO_2_. Distinct colonies, if present, were speciated by MALDI-TOF MS (MALDI Biotyper Sirius CA System, Billerica, MA). Plates without growth were held for additional 7 days before being considered negative. Plates were scored 0 to 4 based on the number of quadrants with bacterial growth.

### Postmortem gross examination & skin histopathology

On day 14 of the ID_50_ study, mice were euthanized with a CO_2_ overdose and the carcass and spleen were weighed, and each animal underwent a complete postmortem gross examination, including macroscopic inspection of the skin and all internal organs. Six mm diameter full thickness skin samples were collected, using a punch biopsy instrument (Integra, Mansfield, MA), at the nuchal crest and approximately 2 cm caudal from the crest in the dorsal lumbar region. Additional samples were taken from lesions, if present. Specimens were fixed in 10% neutral buffered formalin, routinely processed in alcohol and xylene, embedded in paraffin, sectioned at 5 μm, and stained with hematoxylin and eosin (H&E). Histologic examination was performed on all skin biopsies to confirm the presence or absence of Cb-related lesions and to determine the magnitude of the histologic changes. Slides were evaluated and lesions recorded by a board-certified veterinary pathologist (ICM) blinded to groups and doses. Samples were semi-quantitatively scored as normal (0), minimal (1), mild (2), moderate (3), or severe (4), based on the intensity of acanthosis, orthokeratotic hyperkeratosis (orthokeratosis), bacteria and inflammation. Specifically, acanthosis refers to an elevated number of keratinocytes and indicates an increase in epidermal thickness resulting from hyperplasia and often hypertrophy of cells of the stratum spinosum. Minimal acanthosis was defined as an increased thickness of up to a 3-cell stratum spinosum layer; mild as 4 to 7 cell layers; moderate as 8 to 10 cell layers; and, severe as greater than 10 cell layers. Orthokeratosis refers to increased thickness of the stratum corneum without retention of keratinocyte nuclei and was mostly identified in a compact to laminated pattern. Minimal orthokeratosis was defined as an increased thickness of up to 50 μm; mild as 100 to 200 μm; moderate as to 200 to 300 μm; and, severe as greater than 300 μm. Bacterial colonies were assessed based on the presence of distinguishable clusters of short bacilli in the stratum corneum or within the lumen of hair follicles. Minimal presence of bacteria was defined as small amounts of scattered bacterial rods with no formation of characteristic clusters; mild as 1-2 clusters; moderate as 3-5 clusters; and, severe as greater than 6 clusters. Inflammation was mostly comprised of a mixture of mononuclear cells and neutrophils, associated both with and without hair follicle rupture. Minimal inflammation was defined as single superficial pustules and/or small numbers of inflammatory cells scattered in the dermis; mild as slightly larger or increased numbers of superficial pustules and/or slightly higher numbers of inflammatory cells scattered in the dermis; moderate as larger or increased numbers of pustules and/or higher numbers of inflammatory cells in the dermis, frequently forming clusters around capillaries and/or adnexal units; and, severe as large and multiple pustules with high numbers of inflammatory cells forming coalescing clusters or sheets in the dermis, often infiltrating the epidermis and/or adnexal epithelia. For all isolates, the average of each of the 4 microscopic characteristics was calculated per inoculum using 2 biopsies per mouse. The “total histopathology score” was calculated by adding the average of all 4 histologi characteristics. On day 60 of the NSG clinical disease study, mice were euthanized with CO_2_ and an external examination was conducted.

### ID_50_ Determination by Reed and Muench, Dragstedt-Behrens, and Spearman-Karber, & Probit regression

The frequency of positive animals based on culture results per dose was determined and used to calculate the ID_50_. For each isolate and strain of mouse used, the ID_50_ was calculated using the skrmbd package obtained from the USDA in R 4.2.1 statistical computing for Windows (https://www.r-project.org/about.html).^28,37^ The Reed and Muench method was used to calculate the ID_50_ using the two-step equation below. The method uses the cumulative sums of infected animals per dose. First, the proportional distance (PD) between the percentage of positive animals above and below the 50% end point is calculated. The ID_50_ is then calculated by taking the inverse log of the proportional distance and the log of the dose in which less than 50% of the mice became infected.^28,37^

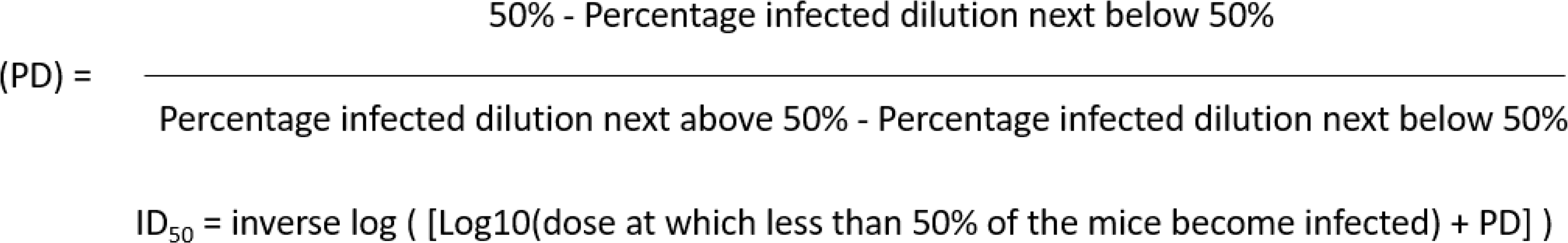

The Dragstedt-Behrens method relies on the frequency of positive animals at each dose and the ID_50_ is estimated by connecting the hypothetical fractions between the doses and determining the intermediate value.^37^ The Spearman-Karber method estimates the 50% end point by the probability mass function of the dose distribution between 0% and 100% response. This method requires at least a single experimental dose to have a 100% positive response to pathogen exposure.^37^ For isolates that did not reach 100% positive responses, the skrmbd package assumes the next lower or higher dose will produce a zero or complete response, respectively. Probit regression analysis was performed using statistical software (MedCalc Software Ltd, Ostend, Belgium).^19^ The probit regression procedures fit a sigmoid dose response curve and calculate values with 95% confidence interval of the dose variable corresponding to a series of possibilities.

### Statistical analysis

The buccal and dorsal culture scores on day 7 and 14 PI were compared using a paired t-test. Differences in percent change of body weight and spleen:body weight ratio, and all histologic characteristics (including hyperkeratosis, acanthosis, bacterial groups, inflammation, and the total histopathology score) were compared for each isolate by dose to their respective control using a one-way ANOVA. Post-hoc testing was performed using Dunnett’s multiple comparison test. *p*-Values ≤0.05 were considered statistically significant. All calculations were performed using Prism 9 for Windows (GraphPad Software version 9.3.0, San Diego, CA).

## RESULTS

### Growth Kinetics of Corynebacterium bovis Isolates

The Cb isolates displayed different growth rates and characteristics (Figure 2). Isolates 7894, 17-0240, Iso 24956, 16-2004, and WCM 35 aggregated forming clumps when grown in broth, whereas the other isolates had uniform turbidity. The bacteria within the clumps were dispersed by gentle manipulation with a pipette prior to densometer readings.

**Figure 1.**
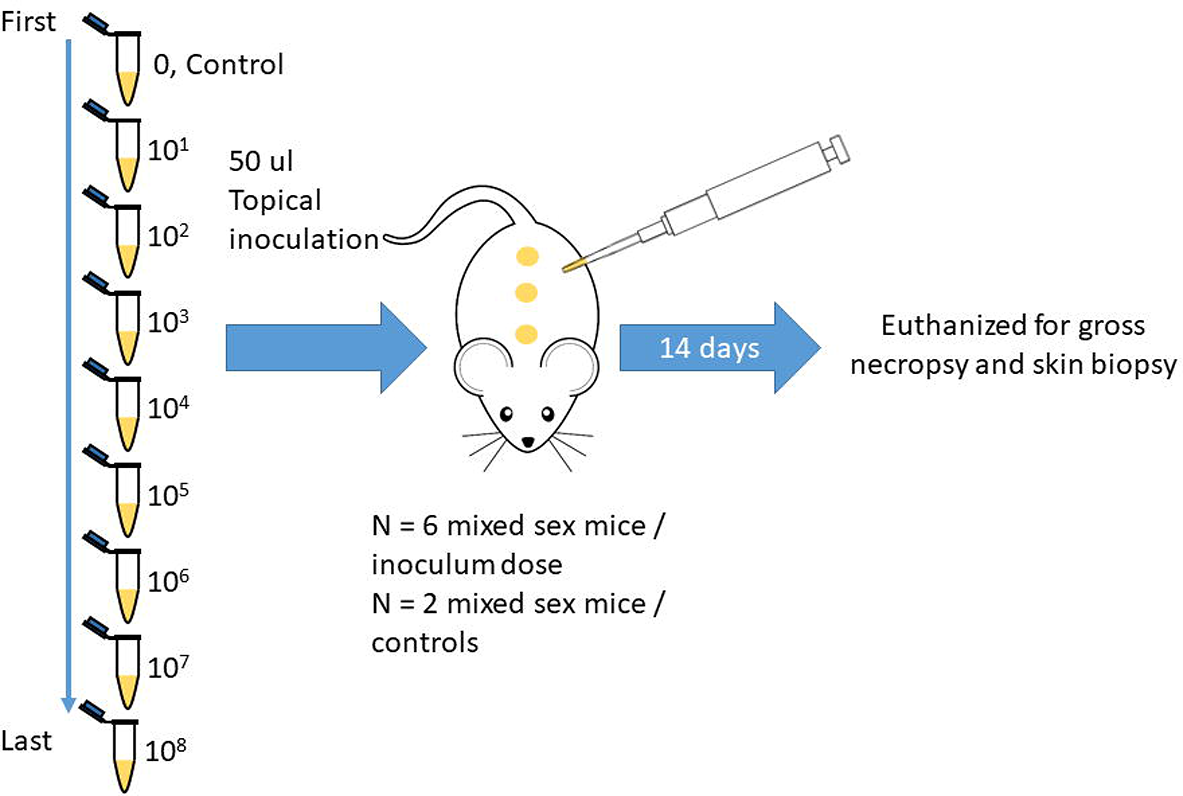
Experimental design: Eight groups of 6 nude mice were inoculated with increasing doses of Cb. The control group consisting of 2 mice received media without bacteria. Animals were monitored daily for clinical signs and cultures (skin and buccal) were conducted weekly until 14 days PI, the study endpoint.

**Figure 2.**
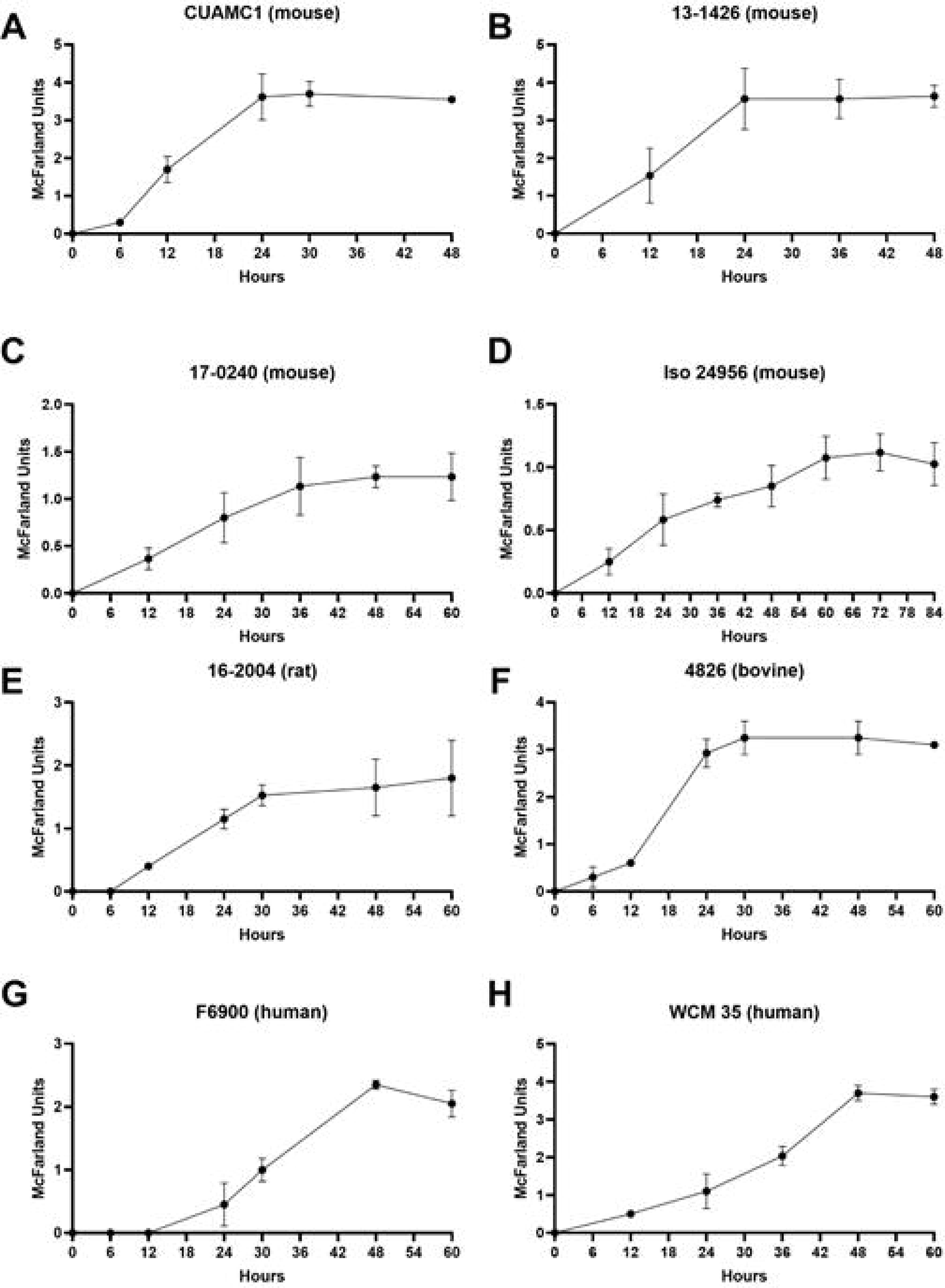
Growth kinetics of *Corynebacterium bovis* isolated from various species. Densometer readings correspond to the average McFarland standard values of three independent experiments and errors bars represent the standard error of the mean (n=3). The growth kinetics of isolate 7894 was previously reported and therefore excluded from the figure. ^10^

The Cb isolates used in this study were slow growing compared to other bacterial species such as *Staphylococcus spp*.^18^ All isolates remained in the lag phase for the first 0 to 12 h followed by an exponential phase that lasted between 12 and 60 h. The mid-log ranged from 13 and 35 h (Table 4). The mouse derived isolates reached mid-log between 13 and 29 h, whereas the bovine and rat isolates reached mid-log at 16 and 20 h, respectively. The human isolates had the slowest growth rates reaching mid-log at 30 or 35 h. The beginning of the stationary phase varied between 24 and 60 h.

**Table 4.** Time to reach mid-log growth for 9 *Corynebacterium bovis* isolates from various species.^10^

| Isolate | Source | Mid-log (h) |
| --- | --- | --- |
| 7894 | Nude mouse | 18 |
| CUAMC1 | Nude mouse | 13 |
| 13-1426 | NSG mouse | 13 |
| 17-0240 | Nude mouse | 20 |
| Iso 24956 | Nude mouse | 29 |
| 16-2004 | Nude rat | 20 |
| 4826 | Bovine | 16 |
| F6900 | Human | 30 |
| WCM 35 | Human | 35 |

### ID_50_ and Dose Response Model

The frequency of mice testing positive by either buccal, dorsal or both culture methods was used to calculate the ID_50,_ and the dose response curves generated by the probit regression analysis. Probit regression models were used to generate dose response curves with 95% confidence intervals (CI) (Figure 3). As isolates 7894, CUAMC1, and 16-2004 in NU mice had groups in which 50% of the mice were colonized, CIs could not be generated. Dose response curves could not be generated for the human isolates, F6900 and WCM 35, nor the NSG mice inoculated with 13-1426 as they remained uninfected following inoculation with as many as 10^8^ bacteria or did not have sufficient positive animals to perform the calculations.

**Figure 3.**
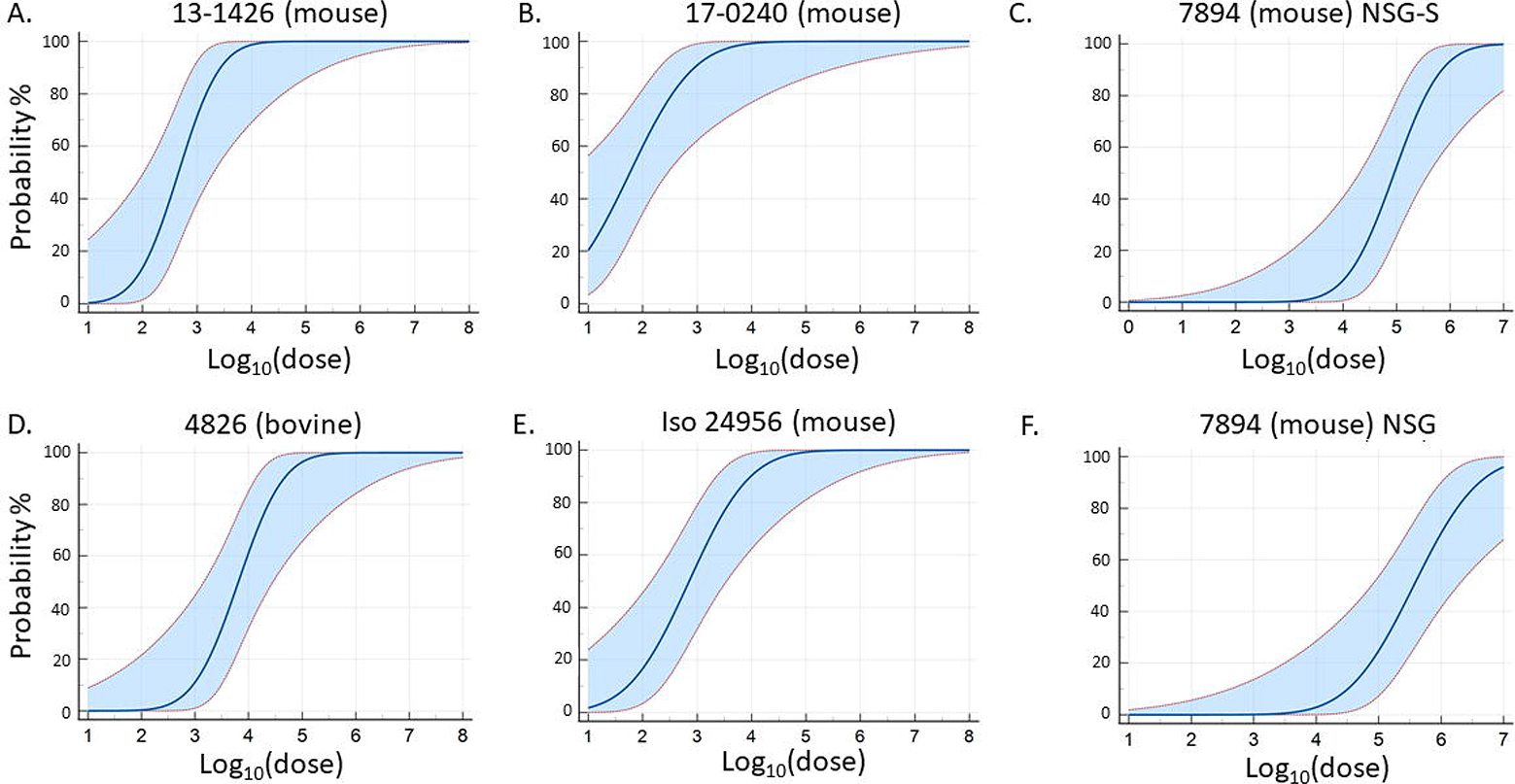
Dose response curves of *Corynebacterium bovis* isolates generated by probit regression analysis. Curves showing the probability of infection and maximum likelihood estimation, ranging from 0 to 100, for different Cb isolates and strain of mouse inoculated indicated in parentheses. Red dashed lines and blue shading depict the 95% confidence interval for dose corresponding to a particular probability.

The ID_50_ for Cb varied by isolate and host species in NU mice (Table 5). Mouse derived isolates had considerably lower ID_50_s (58 to 1,000 bacteria) compared to those obtained from other species. Isolate 17-0240 in NU mice was the only isolate whose probit curve indicated a higher probability (20%) of infection starting at lower doses. Furthermore, 17-0240’s ID_50_ was 100 bacteria when using the Reed and Muench and Dragstedt-Behrens methods but was almost half when determined using probit analysis. The rat (16-2004) and bovine (4826) isolates had ID_50_s between 6,000 to 10,000 bacteria, 60 to 100 times greater than mouse isolates 7894 and 17-0240, which had the lowest ID_50_. No mice were colonized with the human isolates (F6900 and WCM 35) at doses as high as 10^8^ bacteria at either 7- or 14-days PI. Therefore, an ID_50_ could not be determined for either isolate.

**Table 5.**
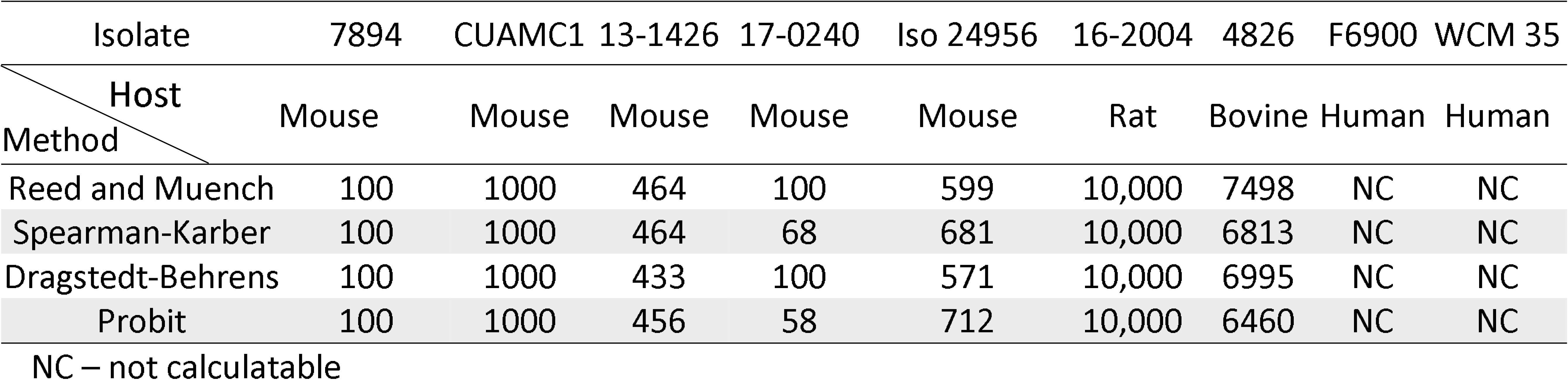
ID_50_ of *Corynebacterium bovis* isolates obtained from different host species in nude mice.

When NSG and NSG-S mice were inoculated with mouse isolate 7894, an ID_50_ between 91,056 to 133,325 and 284,000 to 363,998 bacteria, respectively, was determined depending on the method used to calculate the ID_50_ (Table 6). These doses were approximately 13,000- and 23,000-fold higher than when the same isolate was administered to the NU mouse. When the ID_50_ was determined for 13-1426 (NSG mouse isolate), the ID_50_ could not be determined in the NSG strain as only 2 of 6 mice inoculated with 10^8^ bacteria cultured positive. Therefore, the ID_50_ was most likely greater than 10^8^, whereas the ID_50_ for this isolate was 433 to 464 bacteria in the NU mouse. Probit analysis revealed a rightward shift of the probability curve in furred mice as compared to NU mice, with the slope increasing starting at an inoculum of 10,000 bacteria.

**Table 6.**
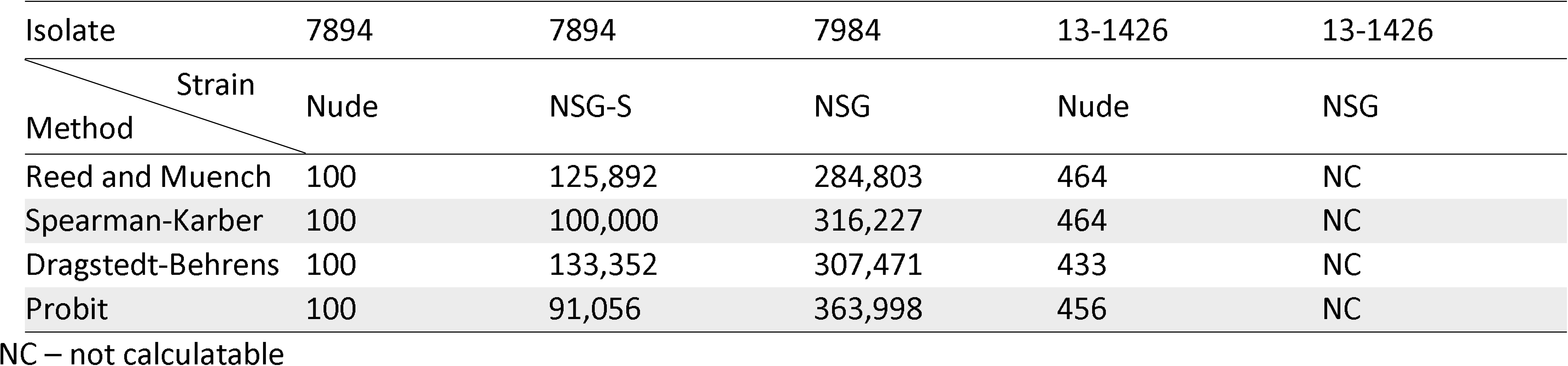
ID_50_ of 2 murine *Corynebacterium bovis* isolates in various immunodeficient strains.

### Disease course in immunocompromised mice

Disease progression and severity differed by Cb isolate in NU mice (Figure 4). If the isolate caused disease, larger inocula generally yielded an earlier disease onset, higher disease scores, and earlier clinical resolution. Animals receiving a Cb isolate that caused moderate to severe disease, e.g., 7894, 13-1426, 16-2004 and 17-0240, depending on the inoculum dose generally reached peak disease and showed or began to show clinical resolution by study termination on day 14 PI (Figure 4). When the isolates that caused clinical disease were compared over the 14-day PI observation period, disease severity was ranked (from most to least severe): 7894, 17-0240, 13-1426, 16-2004, CUAMC1, and 4826, when inoculated with 10,000 bacteria (Figure 5A). CUAMC1 (mouse) and 4826 (bovine) produced minimal disease with only a single animal per isolate developing disease with a score of 2 over the 14-day PI observation period, and this only occurred in mice receiving the highest inoculum of 10^6^ and 10^8^ bacteria, respectively (Figure 4). When the disease scores were plotted by the ID_50_ inocula, or the dose just above the calculated ID_50_ when the ID_50_ did not correspond to the inoculum administered, as well as the lowest dose resulting in clinical signs over the 14-day observation period, disease progression was similar among isolates (Figure 5B and C). Interestingly, despite clinical resolution, some mice receiving higher inocula remained culture positive with a large number of bacteria isolated on culture (culture grade 3-4). Despite some animals testing positive by buccal and dorsal culture, Iso 24956 did not produce disease in NU mice. The human derived isolates F6900 and WCM 35 failed to colonize mice, therefore no animals inoculated with these isolates displayed clinical signs. Although disease severity was not associated with lower mid-log growth rates, mice inoculated with isolates with lower ID_50_s, with the exception of Iso 24956, tended to develop more severe clinical disease, e.g., 7894, 17-0240, and 13-1426.

**Figure 4.**
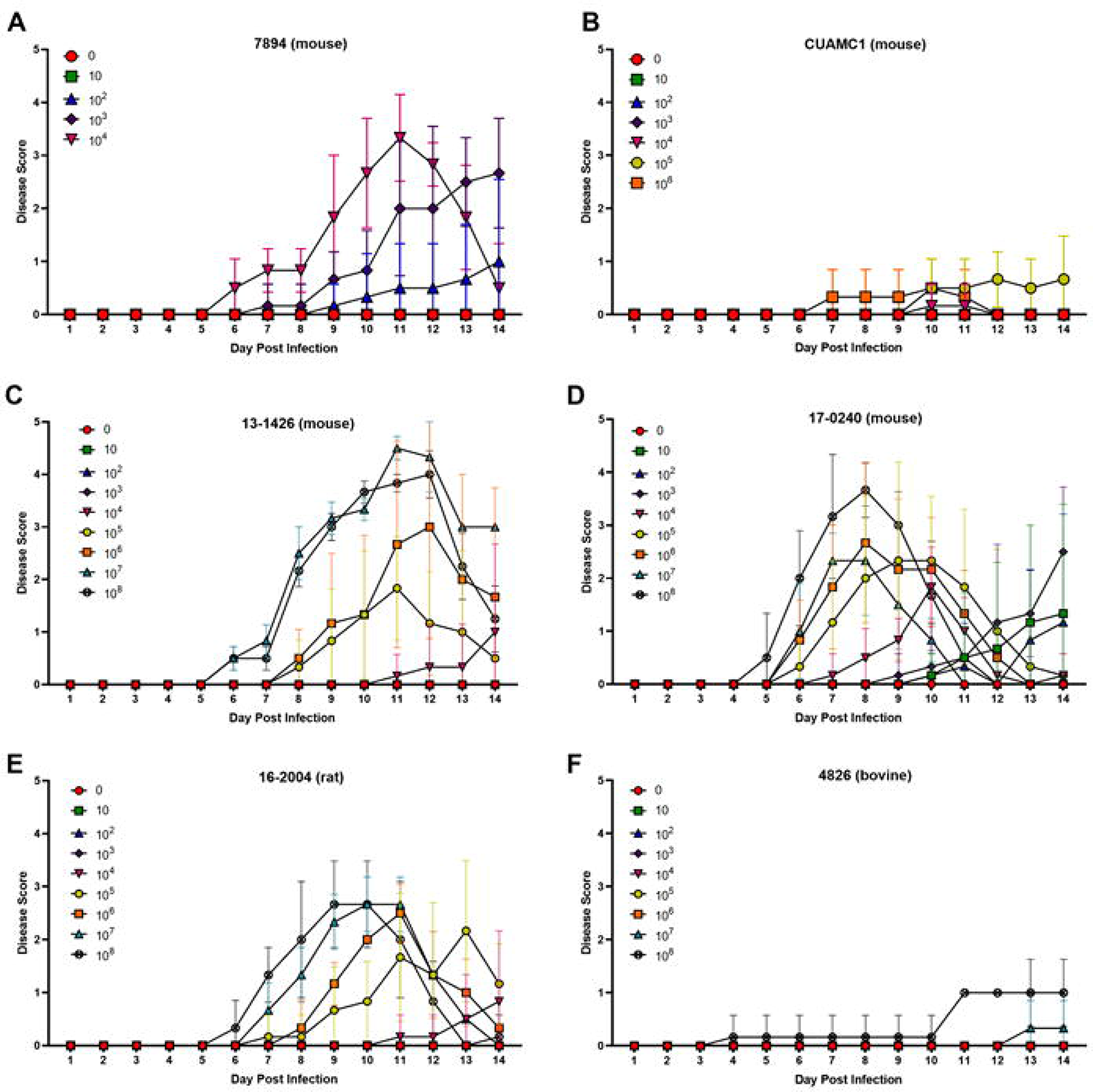
Disease scores by dose for athymic nude mice inoculated with various *Corynebacterium bovis* isolates. Mice were inoculated with 0 to 10^8^ bacteria with the exception of isolates 7894, F6900, and CUAMC1 which received a maximum inoculum of 10^4^, 10 ^6^, and 10 ^6^, respectively. The animals were monitored and scored daily for 14 days. Scores represent the mean +/− standard deviation of 6 mice receiving the same inoculum.

**Figure 5.**
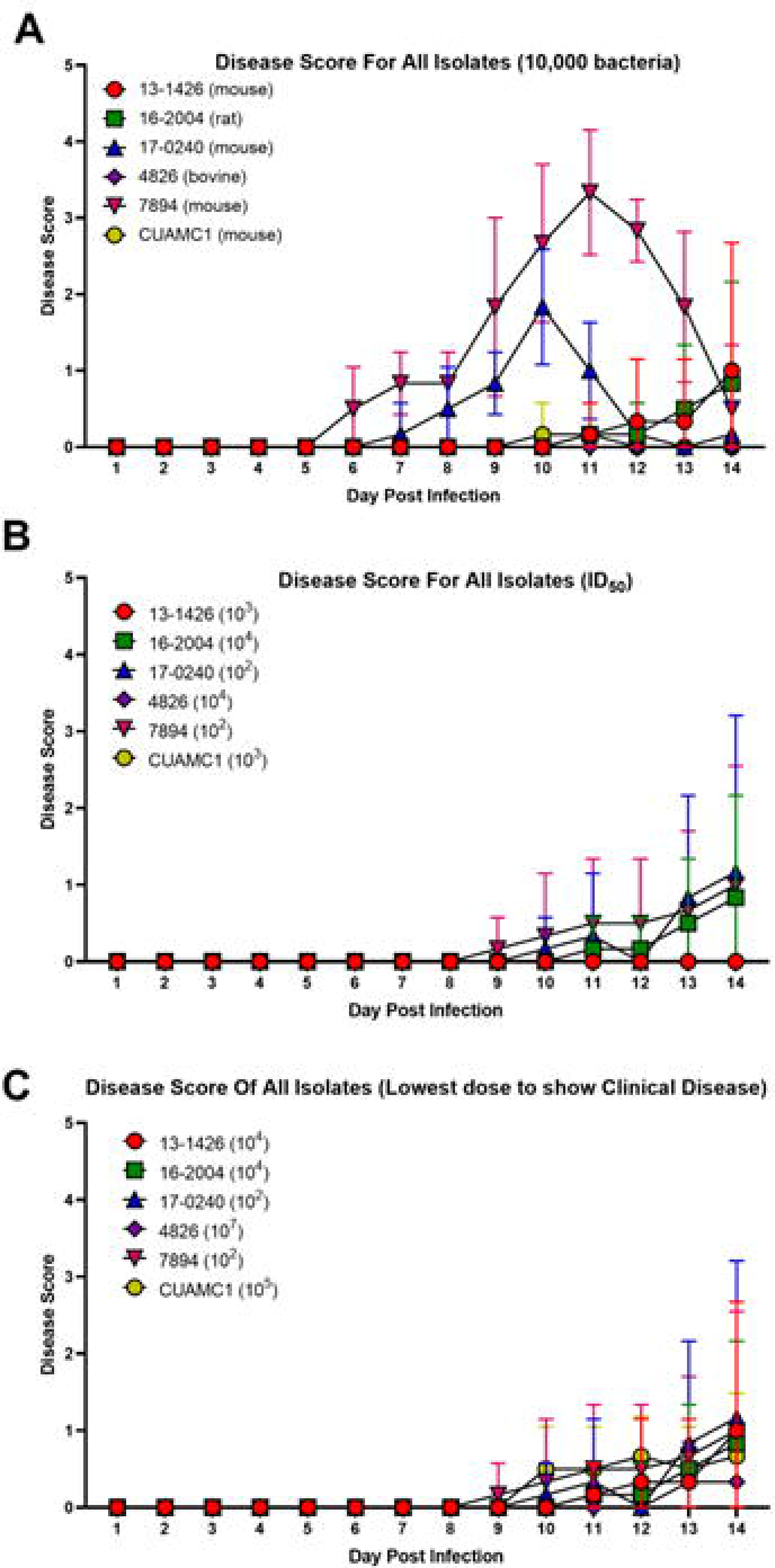
Daily disease scores for 14 days PI for all *Corynebacterium bovis* isolates in athymic nude mice infected with 10,000 bacteria (A), by ID_50_ (B) and the lowest dose resulting in clinical disease (C). Mice were inoculated with 0 to 10^8^ bacteria with the exception of isolates 7894, F6900, and CUAMC1, which received a maximum inoculum of 10^4^, 10^6^, and 10^6^, respectively. Scores represent the mean +/− standard deviation of 6 mice receiving the same inoculum.

**Figure 6.**
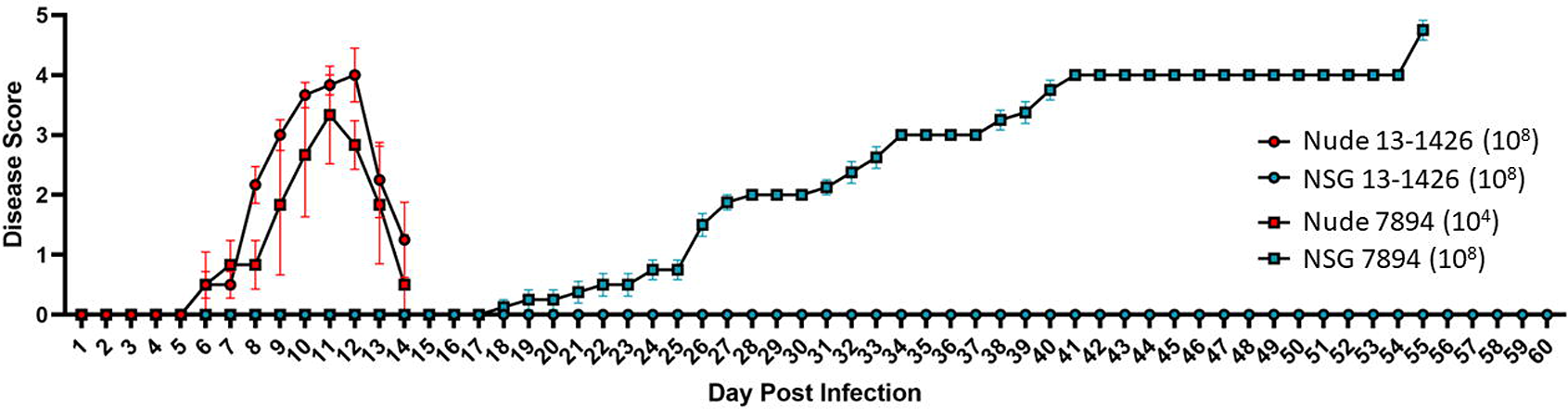
Daily disease scores for 60 days PI for NSG mice infected with *Corynebacterium bovis* 7894 and 13-1426 with 10^8^ bacteria. Scores represent the mean +/− standard deviation of 6 mice receiving the same inoculum.

When buccal and dorsal culture sites were compared for their ability to identify Cb colonized mice, there were no statistical differences noted (data not shown). Culture positive NSG and NSG-S mice did not develop clinical signs over the 14-day observation period when inoculated with either mouse isolate, 7894 or 13-1426. Additionally, culture positive furred immunodeficient mice consistently had fewer bacteria recovered (culture grade 1-3) as compared to culture positive NU mice (culture grade 3-4) at this time point. For the NSG-S strain, larger numbers of bacteria were isolated at day 14 PI (culture grade, with one animal scoring a 4 on buccal culture) as compared to day 7 PI (culture grade 1-2); however, this was not consistently observed with NSG mice where mice at day 7 and 14 PI consistently scored low (grade 1-2 with only 2 animals scoring 2) with only a few colonies present. In the 60-day NSG mouse study, when inoculated with 10^8^ isolate 7894 bacteria, NSG mice presented with clinical disease on day 18 or 22 PI, depending on the cage. All mice in both cages, which included the 4 contact sentinels, developed clinical disease by day 26 and reached the clinical end point (disease score of 5) by day 55. NSG mice inoculated with 10^8^ isolate 13-1426 bacteria remained culture negative and did not develop clinical disease over the 60-day observation period, despite this isolate having been cultured from clinically affected NSG mice.

### Postmortem gross examination & histopathology

Gross lesions, if present at euthanasia, were limited to the skin. Macroscopic skin lesions in NU mice, when present at necropsy on day 14 PI, matched the clinical findings and included varying degrees of white flaking and erythema of the skin over the entire carcass, but especially affecting the ventrum, muzzle, cheeks, and limbs (Figure 7). There were no gross lesions in NSG and NSG-S inoculated with mouse derived isolates 7894 and 13-1426 over the 14-day observation period. However, histologic lesions were noted in some animals infected with 7894 at day 14 (Figure 8). In the 60-day study, at necropsy the NSG mice inoculated with isolate 7894 displayed varying degrees of periorbital swelling, erythema, and/or alopecia, especially affecting the face, ears, and trunk. Gross lesions were absent in NSG mice inoculated with isolate 13-1426.

**Figure 7.**
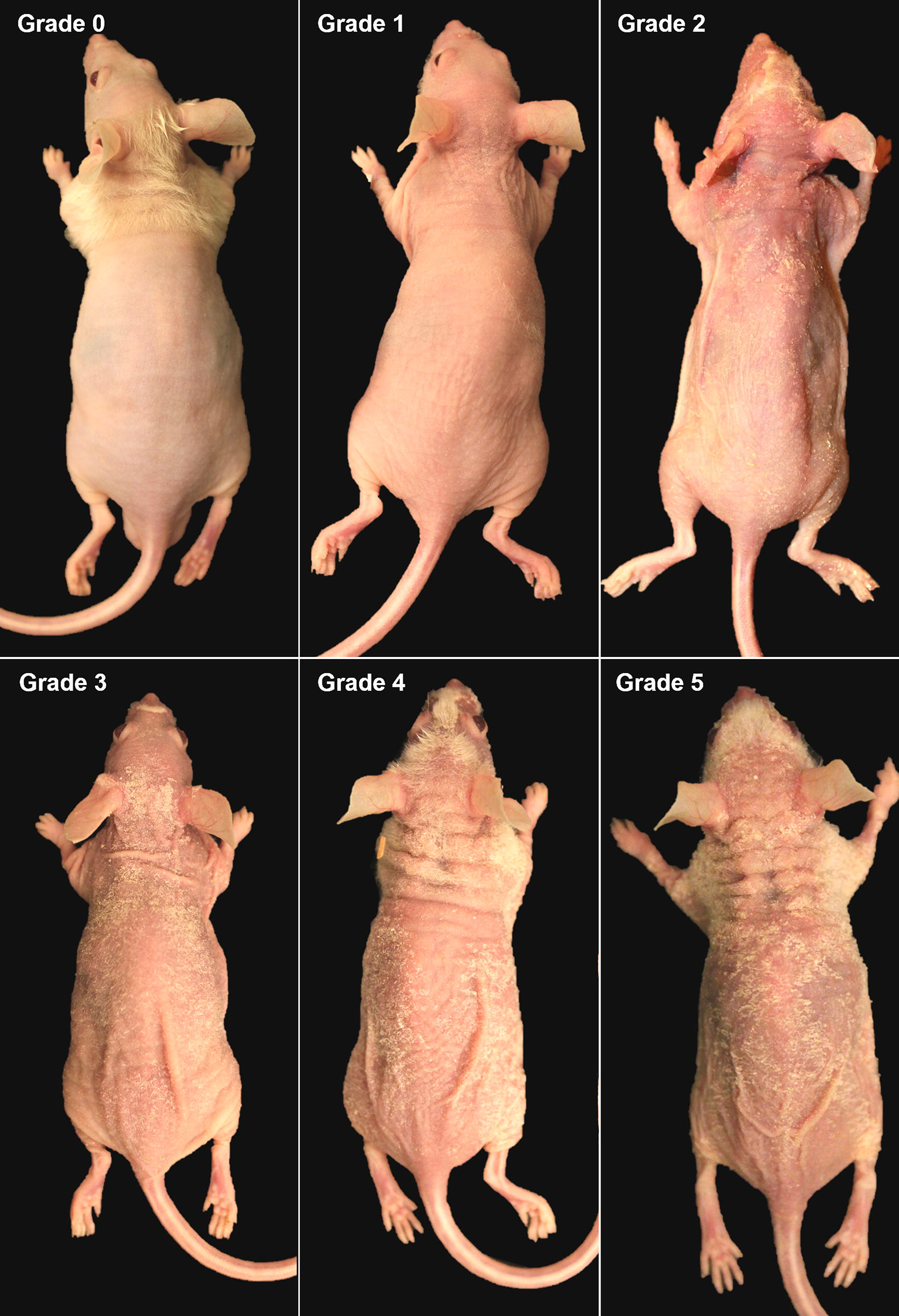
Clinical disease by score at gross necropsy for athymic nude mice. Animals were graded 0 to 5 based on clinical signs representing mild, moderate, and severe disease.

**Figure 8.**
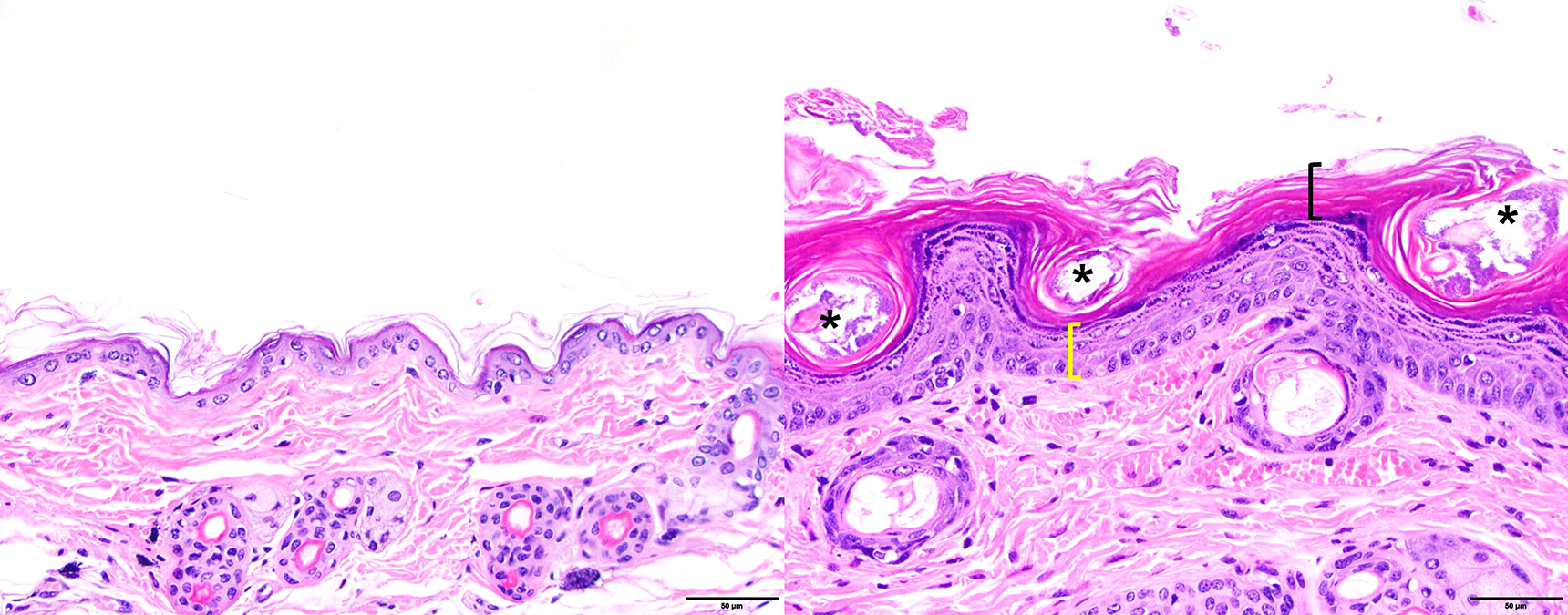
Skin histology of NSG mice infected with Cb 7894 at 10^7^ bacteria at day 14 PI. The left panel depicts normal skin histology. The right panel depicts histologic changes associated with Cb infection. *, bacteria; black bracket, hyperkeratosis; yellow bracket, acanthosis.

There were no significant differences in either the percent change of body weight or the spleen:body weight ratio from any isolate administered at any dose to NU, NSG, and NSG-S mice when compared to the corresponding controls inoculated with bacteria-free media with the exception of NU mice inoculated with ≥ 10^6^ isolate 13-1426 bacteria. The body weight of NU mice inoculated with 10^6^, 10^7^, and 10^8^ isolate 13-426 bacteria had a significant decrease (p = 0.02) of 2.13, 3.76, and 0.59%, respectively, compared to the control group.

On histology, bacteria were observed in 39 of 528 mice receiving any Cb isolate at any dose. If bacteria were present, the majority of animals inoculated with Cb isolates 7894, 13-1426, and 17-0240 were at peak disease (Table 7 and Figures 4). Bacterial colonies were only observed in NSG (10^4^ and 10^7^), and NSG-S (>10^6^) mice infected with high doses of isolate 7894 (Table 8). There were no bacteria observed microscopically in NU mice infected with CUAMC1 (mouse) and F6900 (human) (Table 7 and 9). In a single animal, small bacterial colonies were observed in WCM35 (human) at a dose of 10^2^ bacteria. These colonies had similar morphology and distribution as identified in other Cb isolates; however, WCM35 colonization was not confirmed as these animals were never culture positive.

**Table 7.**
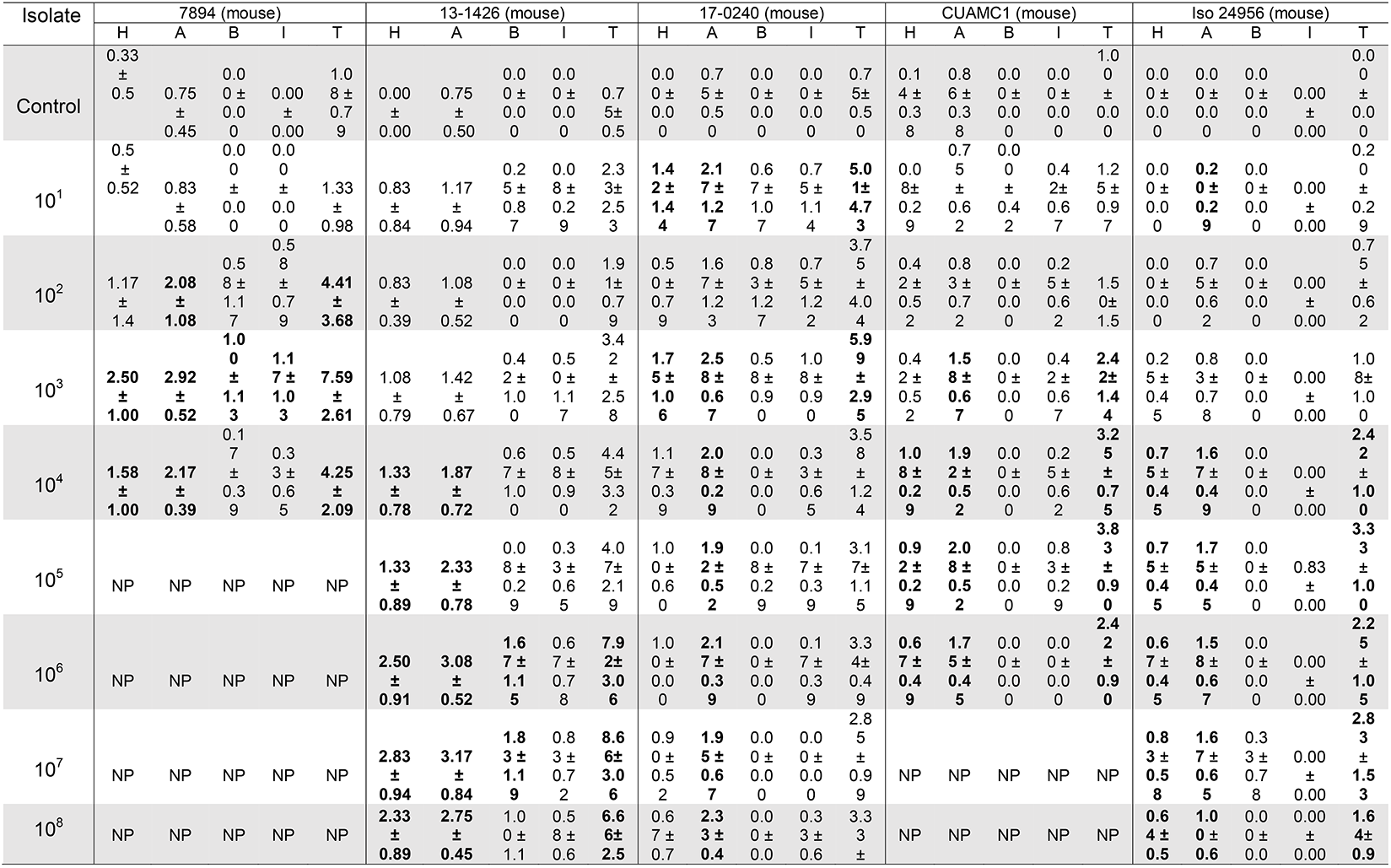
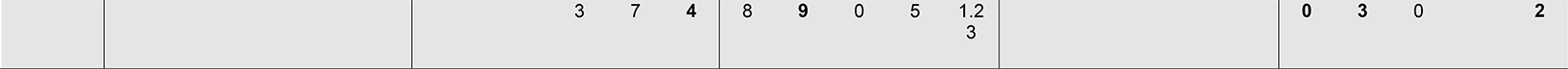
Histopathology scores for athymic nude mice infected with *Corynebacterium bovis* isolates sourced mice. Scores (0 – 5) represent the mean ± standard deviation of 2 biopsies per mouse from a total of 6 mice receiving the same inoculum. Bolded scores are statistically significant compared to their respective controls (p ≤ 0.05). Abbreviations: H, hyperkeratosis; A, acanthosis; B, bacteria; I, inflammation; T, Cumulative histology score (H+A+B+I=T); NP, not performed.

**Table 8.**
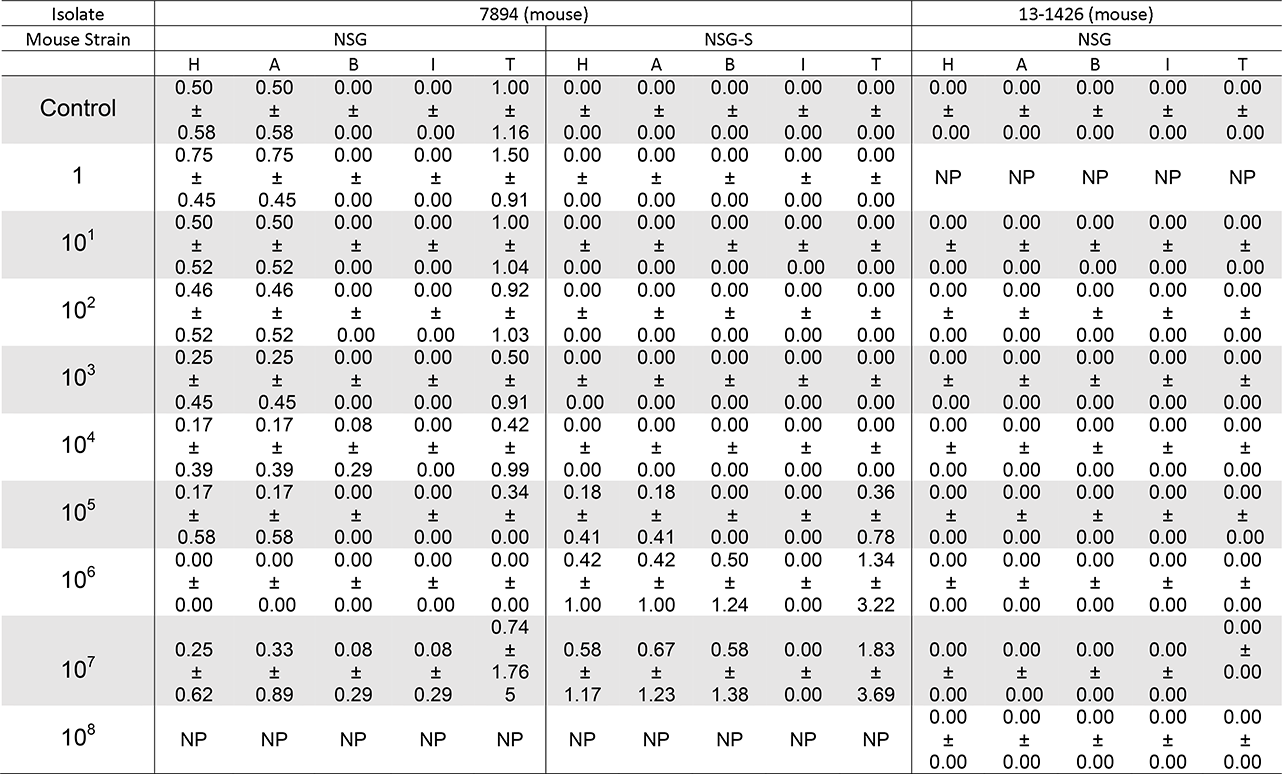
Histopathology scores for haired immunodeficient mice infected with *Corynebacterium bovis* isolates 7894 or 13-1426. Scores (0 - 5) represent the mean ± standard deviation of 2 biopsies per mouse from a total of 6 mice receiving the same inoculum. Bolded scores are statistically significant compared to their respective controls (p ≤ 0.05). Abbreviations: H, hyperkeratosis; A, acanthosis; B, bacteria; I, inflammation; T, Cumulative histology score (H+A+B+I=T); NP, not performed.

**Table 9.**
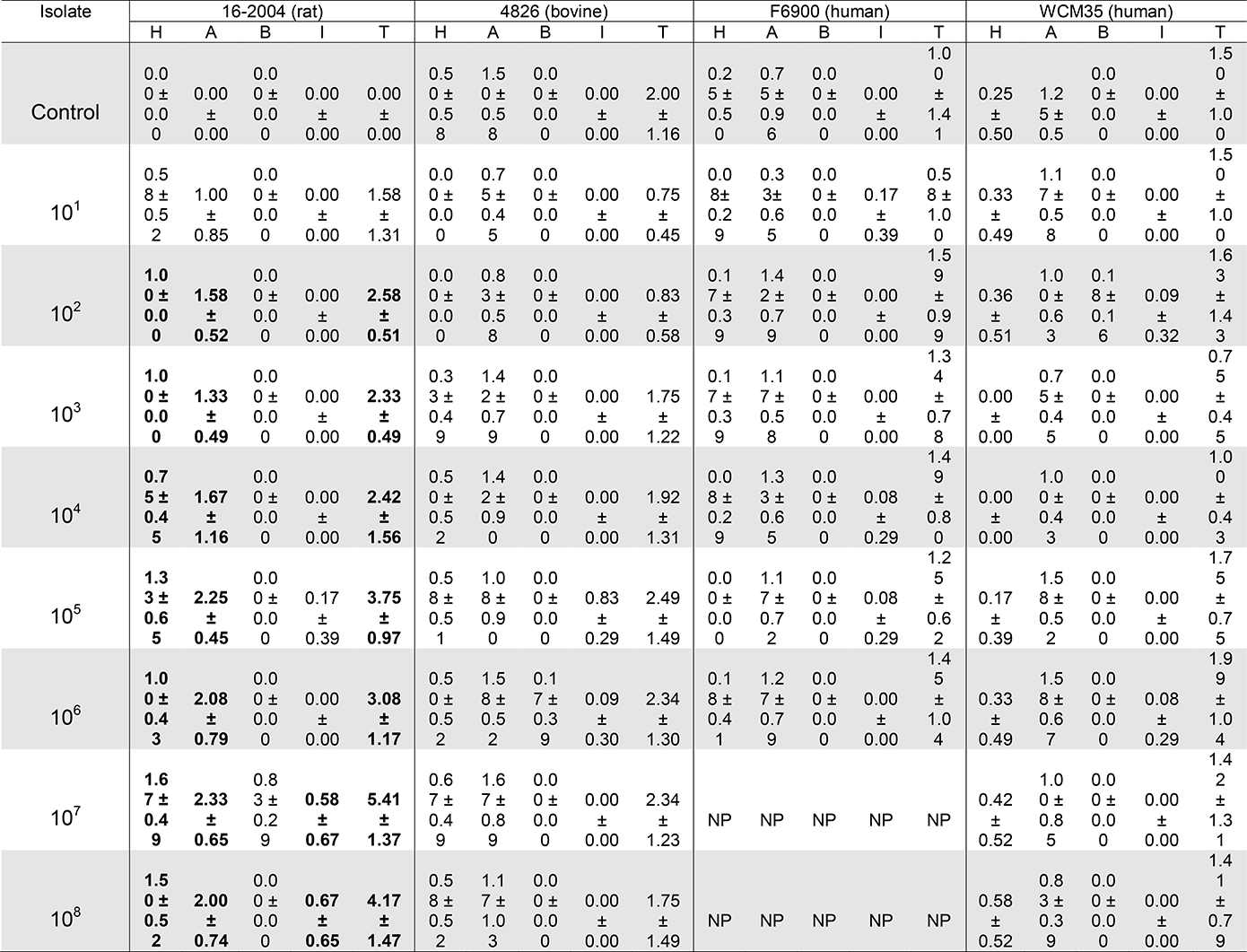
Histopathology scores for athymic nude mice infected with *Corynebacterium bovis* isolates sourced from other species. Scores (0 - 5) represent the mean ± standard deviation of 2 biopsies per mouse from a total of 6 mice receiving the same inoculum. Bolded scores are statistically significant compared to their respective controls (p ≤ 0.05). Abbreviations: H, hyperkeratosis; A, acanthosis; B, bacteria; I, inflammation; T, Cumulative histology score (H+A+B+I=T); NP, not performed.

Dermatitis was observed in NU mice inoculated with select doses of mouse isolates 7894, 13-1426, 17-0240, and CUAMC1, as well as controls (Table 7). When present, it was most often characterized by mixed mononuclear and neutrophilic infiltrates primarily associated with hair follicle rupture and not necessarily Cb infection as was similarly observed in the uninfected controls. The remaining isolates rarely had inflammatory lesions. Inflammation was not observed in any of the NSG nor NSG-S mice examined (Table 8).

NU mice inoculated with mouse Cb isolates had the highest total histology scores primarily consisting of increased scores in acanthosis and hyperkeratosis (Table 7, Figure 9). For all isolates that caused disease in NU mice, the highest total histology scores were associated with higher clinical disease scores at the time of euthanasia and not necessarily higher doses of inoculum (Table 7, Table 9, and Figure 4). The highest total histology scores were observed in mice inoculated with isolates 7894 (7.59), 13-1426 (8.66), and 17-0240 (5.99) (Table 7). For Cb isolates 7894 and 13-1426, the highest scores, 7.59 and 8.66, were observed in mice receiving 10^3^ and 10^7^ bacteria, respectively (Table 7). When compared to the NU mice, the lesions observed in all NSG and NSG-S mice inoculated with Cb were significantly lower than the NU mouse infected with the same isolate (p <0.0001). The NSG and NSG-S mice infected with isolate 7894 (mouse) had histologic scores similar to the control (Table 8). NSG mice infected with 13-1426 did not have any histologic lesions and therefore had a mean score of 0. Regardless of dose, Iso 24956 (mouse) had the lowest total histopathologic scores of the mouse isolates able to colonize NU mice (Table 7). Interestingly, despite not displaying overt clinical disease, NU mice inoculated with Iso 24956 (mouse) had significantly higher mean total scores at doses >10^4^ as compared to the control. Similarly, mouse isolates 7894, 13-1426, and CUAMC1 had significantly higher mean total scores as compared to controls at doses ≥10^2^, 10^6^, and 10^3^, respectively.

**Figure 9.**
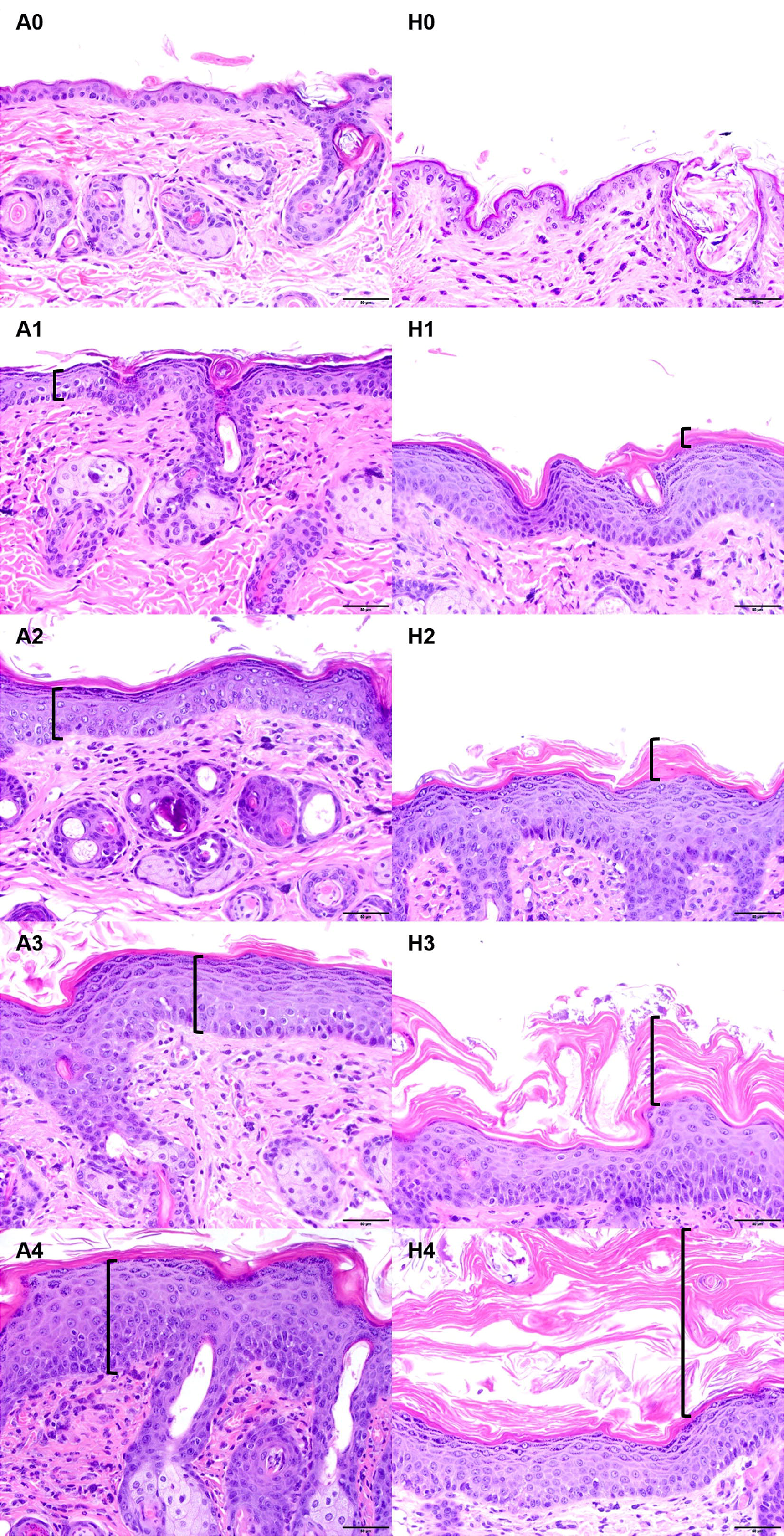
Histologic scoring system for acanthosis and hyperkeratosis in athymic nude mice infected with Cb at day 14 PI. Samples were semi-quantitatively scored as normal (0), minimal (1), mild (2), moderate (3), or severe (4), based on the intensity of acanthosis and orthokeratotic hyperkeratosis (orthokeratosis). A 0-4, acanthosis grade; H 0-4, hyperkeratosis grade; Black bracket represents the amount location and amount of acanthosis (A) or hyperkeratosis (H).

Isolates cultured from humans and cows (WCM35 [human], F6900 [human] and 4826 [bovine]) produced minimal histologic changes associated with Cb colonization despite being inoculated with as many as 10^8^ bacteria (Table 9). When comparing only Cb isolates obtained from other species, NU mice inoculated with rat isolate 16-2004 had the most significantly different hyperkeratosis and total histopathologic scores as compared to the control at doses ≥ 10^2^ (Table 9).

## DISCUSSION

Cb has and continues to pose an extraordinary challenge to institutions using immunocompromised mice as models in various scientific disciplines. Once introduced into a naïve colony, the bacterium is extremely challenging to eradicate.^24^ Despite Cb being first described over 40 years ago, relatively little is known about the bacterium. This study provides striking new and important insights into Cb’s biology. We recently reported genetic differences among Cb isolates from various species, however it was unclear as to whether these differences translated to differences in an isolate’s virulence, i.e., its’ ability to colonize a host, infect and transmit between species and cause clinical disease.^9^ In this study, we have demonstrated that there are considerable differences in all these characteristics among isolates, both those cultured from the same as well as different species.

In public health, quantitative microbial risk assessment is used to determine the overall risk a potential pathogen poses to the public, particularly for water or food borne illnesses.^19,37^ The assessment involves hazard identification, exposure route characterization, dose response analysis, risk evaluation, and management.^19^ While some of these characteristics of Cb have been determined and used to develop strategies to prevent the initial introduction of the bacterium, an important consideration in understanding the infectivity of a microorganism is to determine the number of microbial units necessary to effectively colonize and infect its’ host, i.e., the infectious dose or the minimum number of organisms required to produce infection in a host and allow re-isolation after the host has had time to eliminate the microbe.^2,7,8^ The infective dose capable of producing infection in 50% (ID_50_) of a population is often determined to help define the risk to a specific population.^7,29^ The ID_50_ of Cb was heretofore unknown, therefore the principal aim of this study was to identify the ID_50_ and develop a dose response curve for Cb in immunodeficient mice and determine the inter-species transmissibility of Cb isolates isolated from various host species. Additionally, we aimed to determine whether there were differences in immunodeficient mouse strain susceptibility to Cb. Three methods, the Reed–Muench, Dragstedt– Behrens, and Spearman–Karber, were used to estimate the ID_50_ from dose response data.^37^ We also used a probit logistic regression model to define the probability of infection based on any dose, not just the ID_50_.^19^

It was surprising that the murine Cb isolates evaluated differed markedly in the number of bacteria needed to colonize a murine host. There was as much as a 10-fold difference in the median dose necessary to colonize an NU mouse among the 5 mouse isolates we evaluated in this study, with isolate 7894 not only requiring one of the lowest inoculums to colonize, but it was also the most virulent in that it caused the most severe clinical disease with the earliest onset when compared to other isolates at the same dose. In contrast, CUAMC1 required 10 times more bacteria to colonize a NU mouse and caused minimal clinical disease. Equally surprising was the finding that neither of the 2 human isolates evaluated resulted in colonization, even though the mice were administered as many as 10^8^ bacteria. While additional human isolates will need to be evaluated, these findings suggest that Cb isolates infecting humans may pose little to no biosecurity risk to laboratory mice.

Also unexpected, was the finding that the ID_50_ of the most virulent and one of the most infectious isolates (based on clinical scores and its ID_50_ in NU mice), mouse isolate 7894, was several magnitudes higher in the furred highly immunocompromised NSG and NSG-S strains as compared to NU mice. Despite the NSG and NSG-S strains being considerably more immunodeficient than NU mice, 1,000 to 3,000 times more Cb needed to be inoculated to achieve colonization and none of the mice developed clinical signs during the 14-day PI observation period. As a result of these findings and our suspicions based on clinical experience that the onset of clinical disease is considerably later in NSG mice, NSG mice were inoculated with either isolate 7894 or 13-1426 and observed for 60 days. NSG mice administered isolate 7894 began to show clinical signs as late as day 21 PI, considerably later than when this isolate was administered to NU mice. However, in contrast to the NU, clinical disease progressed until they reached humane endpoint at day 55 PI in the NSG strain. A recent study described requiring 10-fold higher doses of CUAMC1 to infect NSG (2×10^7^ cfu) as compared to NU mice (2×10^6^ cfu) and none of the mice developed clinical disease during the 70-day PI observation period.^25^ Despite these contrasts in median infectious dose among these commonly used immunocompromised strains, infection and the resulting clinical disease remains high in enzootically infected vivaria using these and other highly immunocompromised strains. One consideration is that natural infection may occur because of multiple exposures to Cb and could be facilitated by altered skin biology and flora resulting from hair removal by depilatory agents or shaving during tumor implantation or surgery. Also of interest was the finding that isolate 13-1426, originally cultured from an NSG mouse with Cb-associated clinical disease, had an ID_50_ greater than 10^8^ in NSG mice. In contrast, less than 500 bacteria were needed to colonize 50% of the NUs. None of the 8 mice inoculated with this isolate and followed for 60 days PI were colonized. While no NSG mice became colonized with this isolate in the 60-day study, 2 of 6 mice did become colonized when 10^8^ bacteria were administered when attempting to define the ID_50_. It would be valuable to understand how long after NSG mice become colonized with this isolate that clinical signs develop.

To determine the ID_50_, an understanding of each isolates’ growth kinetics was necessary as bacteria needed to be inoculated when their growth was at mid-log, a time when they are actively dividing. All Cb isolates grew slowly, which was consistent with our previous findings for this species and is the reason why cultures should be held for 5 to 7 days before ascertaining that the organism is absent in samples.^10^ The majority of isolates tended to cluster together, with most mice and human isolates having similar mid-log growth times (13 to 20 and 30 to 35 hours, respectively). Although collectively the isolates growth was slow, isolates grew at different rates with only the mouse isolates, CUAMC1 ,13-1426, and 17-0240, and the rat isolate 16-2004 having similar mid-log growth times of 13 and 20 h, respectively. However, a shorter time interval to mid-log growth was not associated with a lower ID_50_, a shorter onset of clinical disease, or greater disease severity. Mouse isolates CUAMC1 and 13-1426 shared similar times to attain mid-log growth, but 13-1426 displayed severe disease as compared to CUAMC1 which displayed minimal disease. Similarly, a longer time interval to reach mid-log growth did not correlate with a lack of colonization as mouse isolate Iso 24956, 1 of the 3 slowest growing Cb isolates, colonized NU mice, but clinical disease did not develop during the PI observation period. However, both human isolates (F6900 and WCM 35), which had the slowest growth rates, failed to colonize NU mice. Whether these isolates growth characteristics played a role in their inability to colonize remains unknown. It is also important to consider that the bacterium’s growth kinetics likely differ *in vitro* and probably vary by isolate and mouse strain colonized.

Isolates 7894, 16-2004, Iso 24956, 17-0240, and WCM 35 developed auto-aggregation in vitro, a feature demonstrated by other *Corynebacterium* species and other bacteria.^33^ The process is mediated by self-recognizing surface structures linking proteins and exopolysaccharides.^33^ The biological function is not fully understood but is thought to protect bacteria from environmental stress or host responses by promoting biofilm formation, increasing tolerance to antimicrobial agents, impeding phagocytosis by the host immune system, or elevating the invasion frequency of the host’s cells.^1,12,17,36^ While Cb isolates such as 7894 produced severe disease and developed aggregates, others that aggregated in vitro such as Iso 24956 resulted in colonization without disease or, as with WCM 35, failed to colonize at all. Thus, auto-aggregation alone cannot be used to predict Cb’s ability to infect or cause disease.

NU mice are overtly susceptible to Cb which accounts for the high morbidity that has been previously reported.^11^ However, when comparing the clinical signs presented by the infected NU mice used in this study, disease varied by isolate and dose. For example, mice colonized with isolates 7894 and 17-0240 displayed signs of CAH when administered 10^3^ bacteria; however, CUAMC1, Iso 24956, and 13-1426 did not. With higher inocula, mice colonized with isolates 7894, 13-1426, and 17-0240 showed significant disease with some animals reaching a clinical score of 5. In contrast, mice colonized with the remaining isolates had mild, minimal, or no disease. Interestingly, mice colonized with CUAMC1 in our study showed minimal disease with only 8 of 30 animals showing clinical score of 1, and an additional animal which scored 2 at day 14 PI. In contrast, in a recently published article in which NU mice were inoculated with this isolate the authors reported there were no obvious clinical signs; however, hyperkeratosis was identified microscopically over the 70-day observation period.^25^ We hypothesize that differing inoculation methods and housing conditions, the phase of growth at which the bacteria were administered (24 h) and the media used (PBS) likely accounted for this difference.^25^

When NU mice developed clinical disease in this study, CAH was observed between 6 to 14 days PI, depending on the isolate and dose, with some animals reaching clinical resolution within the 14-day PI period. These results were similar to those in previous reports.^11,13,30^ While a virulent Cb isolate obtained from a NU mouse with severe CAH has been described, when NU mice were inoculated with this isolate and compared to mice inoculated with a less virulent isolate also cultured from NU mice, as well as the bovine ATCC isolate, no differences in clinical disease were noted as all isolates caused mild CAH.^13^ The authors speculated that this finding may have resulted from attenuation of the virulent isolate when it was grown *in vitro*. The cause of differing CAH presentations in NU mice is hypothesized to be multifactorial, with experimental manipulation, skin flora, humidity, type of caging, and hair growth cycles impacting the clinical phenotype.^5,11,25^ Herein, we clearly demonstrated that the resulting clinical phenotype varies depending on the isolate with which the mouse is colonized. We could not identify specific putative virulence factors contributing to the observed differences, as many of the previously identified factors of interest were shared among isolates causing markedly different clinical phenotypes.^9^ Further studies will be needed to identify what contributes to Cb’s pathogenicity.^34^

The inter-species transmissibility of bovine adapted Cb has previously been reported.^13^ To our knowledge this is the first study evaluating the potential of Cb isolated from humans and rats to infect immunocompromised mice. Compared to Cb isolated from mice, the ID_50_s of Cb isolated from the rat and cow are as much as 10-times higher. Even though their ID_50_s are higher, the potential remains for these species to act as reservoirs and therefore appropriate biosecurity precautions should be taken. Caution should be exercised when housing nude rats in the same room as immunocompromised mice. Moderate to severe clinical disease was noted in NU mice colonized with the rat Cb isolate 16-2004, while the rat from which this isolate was cultured had ulcerative lesions on its limbs, dorsum, and paws it was unclear if these were due to an active Cb infection or the result of other bacteria isolated which included *Staphylococcus aureus* and *Staphylococcus xylosus*. It is unknown if nude rats will develop Cb-associated disease, or if they will remain colonized with Cb. NU mice can be colonized with a bovine isolate as had been shown previously; the isolate we evaluated produced mild disease and only at the highest doses of 10^7^-10^8^ bacteria.^13^ The isolates cultured from humans failed to colonize NU mice at doses as high as 10^8^ bacteria. These isolates were cultured from the eye and a skin wound and had the longest growth times. It is unlikely that humans act as a reservoir for isolates capable of causing Cb-associated disease in immunocompromised mice. However, Cb (presumably of mouse origin) was isolated from the nasal passages of an animal care staff member working extensively with enzootically infected immunocompromised mice.^6^ Therefore, it is likely that humans could minimally serve as a fomite in the transmission of Cb.

Isolate Iso 24956, which colonized NU mice but did not cause disease, is notable. This isolate was cultured from a commercial breeding colony of highly immunocompromised mice which never developed clinical signs. Subsequent studies, conducted by the breeder, in which various hairless and furred immunocompromised and immunocompetent mice were inoculated with the isolate failed to demonstrate clinical disease despite some of the strains becoming colonized. Whole genome sequencing of the isolate revealed it clustered with that of the previously sequenced cow and human isolates, but it was closest to the sequenced human isolates. While the isolate’s ID_50_ was closest to the rodent isolates, its growth kinetics were similar to that found in the human isolates. The commercial breeder considered the possibility that the isolate was introduced into the affected barrier by a staff member, some of which do have contact with cows, however the definitive source of this isolate remains unknown. The question remains as to whether human isolates given a sufficient dose and/or with prolonged exposure could adapt to colonize the mouse’s skin and cause disease. Future studies with additional human isolates, higher inocula, and/or repetitive administrations should be undertaken to confirm our preliminary findings.

Hairlessness, the magnitude of immunosuppression, and altered skin homeostasis are thought to contribute to the mouse’s susceptibility to Cb, as hairless immunocompetent SKH1-Hr^hr^ and epidermal-mutant *dep/dep* mice were shown to become colonized with the bacterium.^11,26,35^ Skin homeostasis is complex and relies on a variety of factors, including the skin’s microbiome, the hair growth cycle, disintegrating sebocytes releasing lipid rich sebum, continuous replication of epithelial cells, and innate antimicrobial peptide expression.^4,16^ Cb is unable to produce its own lipids, therefore, to survive the organism resides in the stratum corneum utilizing the lipid rich sebum for growth.^32^ Persistent colonization of the immunocompetent *dep/dep* strain with Cb is thought to be mediated by excessive sebum production from hyperplastic sebaceous glands and an abundant amount of lipids in the interfollicular epidermis.^26^ To our knowledge, differences in the amount of and function of sebaceous glandular tissue nor sebum production between NU and furred immunocompromised mice have not been investigated. However, previous studies have shown that Cb grows more rapidly and in greater numbers on the skin of NU compared to NSG mice.^25^ In this study, the mouse’s fur was parted and Cb was inoculated directly onto the skin, therefore the higher ID_50_s determined in the furred immunocompromised strains is likely associated with innate skin defenses such as an increased surface area for antimicrobial peptide accumulation, less sebum production, or differences in the skin microbiome.^25^ Interestingly, the NSG-S strain appeared to be more susceptible to infection as compared to the NSG strain based on the NSG-s’s lower ID_50_ and slightly higher histologic scores when inoculated with isolate 7894 at doses ≥10^6^.

The most significant histologic changes noted in NU mice were associated with rodent isolates, including mouse isolates 7894, 13-1426, and 17-0240, and rat isolate 16-2004. Hyperkeratosis and acanthosis were the major contributors to the total histology scores for all isolates in this study (Table 7–9). Bacteria was infrequently noted despite some animals having severe clinical disease. Histology is a relatively insensitive method to detect pathogens and the absence of bacteria in the examined sections does not rule out their direct involvement in the histologic lesions. Several control mice had hyperkeratosis, acanthosis, and inflammation, which were attributed to the genetic mutation associated with the NU phenotype causing dysfunctional hair development and differentiation.^15,21,27^ Consequently, the inflammation observed in this study was principally associated with folliculitis and rupture of hair follicles, which are spontaneous background lesions in this strain secondary to their dysfunctional follicular structures, and not overtly associated with Cb infection. Interestingly, Iso 24956 resulted in significantly higher total histology scores for doses greater than 10^4^ compared to the control despite animals not showing any clinical signs suggesting that Cb isolates that do not cause overt disease may still alter the skin. Lastly, cross isolate histologic comparisons were not possible because clinical disease varied at the study end point with some animals having higher clinical scores in the lower dose groups as compared to higher doses where clinical signs had begun to regress or had completely resolved.

In conclusion, we have demonstrated considerable differences in individual Cb isolates’ ID_50_s and their ability to cause disease, and if disease developed, its severity. Surprisingly, severely immunocompromised mice require higher inocula to become colonized and clinical disease develops later as compared with NU mice infected with the same isolate. NU mice could not be colonized with Cb isolated from humans with Cb-associated disease. At least some bovine and rat isolates are infectious to mice, although higher doses are needed to achieve colonization. We were unable to identify any genetic or in vitro growth characteristics that could be attributed to any of our findings. Future studies are necessary to elucidate the etiology(ies) of these differences focusing on those that can be harnessed to eradicate Cb from enzootically infected colonies or mitigate its impact.

## Abbreviations

Cb: *Corynebacterium bovis*
CAH: *Corynebacterium*-associated hyperkeratosis
ID_50_: infectious dose 50

## ACKNOWLEDGMENTS

The authors would like to thank Irina Dobtis, King Chan, Morganne Campbell, Noah Mishkin, Michael Palillo and Jackie Candelier for their help and expertise with NU mouse colony maintenance, animal testing and data collection. Additionally, we would like to thank Christopher Manuel and Lars Westblade for providing Cb isolates for this study, and Amine Alioua, Jim Fahey and Anthony Mourino for sharing their Cb isolate, details of their investigations conducted utilizing this isolate, and their perspectives on Cb’s pathogenesis. This research study was funded in part through the NIH/NCI Cancer Center Support Grant P30-CA008748 through Memorial Sloan Kettering Cancer Center.

